# TNF Superfamily Member 14 Drives Post-Influenza Depletion of Alveolar Macrophages Enabling Secondary Pneumococcal Pneumonia

**DOI:** 10.1101/2024.07.28.605445

**Authors:** Christina Malainou, Christin Peteranderl, Maximiliano Ruben Ferrero, Ana Ivonne Vazquez-Armendariz, Ioannis Alexopoulos, Julian Better, Mohammad Estiri, Hendrik Schultheis, Judith Hoppe, Maria-Luisa del Rio, Jose Ignacio Rodriguez-Barbosa, Klaus Pfeffer, Stefan Günther, Mario Looso, Achim D. Gruber, István Vadász, Ulrich Matt, Susanne Herold

## Abstract

Secondary bacterial infection, often caused by *Streptococcus pneumoniae* (Spn), is one of the most frequent and severe complications of influenza A virus (IAV)-induced pneumonia. Phenotyping of the pulmonary innate immune landscape after IAV infection revealed a significant depletion of the tissue-resident alveolar macrophage (TR-AM) population at day 7, which was associated with increased susceptibility to Spn outgrowth. To elucidate the molecular mechanisms underlying TR-AM depletion, and to define putative targets for treatment, we combined single-cell transcriptomics and cell-specific PCR profiling in an unbiased manner, using *in vivo* models of IAV infection and IAV/Spn co-infection. The TNF superfamily 14 (TNFSF14) ligand-receptor axis was revealed as the driving force behind post-influenza TR-AM death during the early infection phase, enabling the transition to pneumococcal pneumonia, while intrapulmonary transfer of genetically modified TR-AMs and antibody-mediated neutralization of specific pathway components alleviated disease severity. With a mainly neutrophilic expression and a high abundance in the bronchoalveolar fluid (BALF) of patients with severe virus-induced ARDS, TNFSF14 emerged as a novel determinant of virus-driven lung injury. Targeting the TNFSF14-mediated intercellular communication network in the virus-infected lung can, therefore, improve host defense, minimizing the risk of subsequent bacterial pneumonia, and ameliorating disease outcome.

## Introduction

Bacterial pneumonia, often due to Spn, is one of the most common complications of primary IAV-induced infection, with a dramatic impact on disease outcome (1). Risk of death, intensive care unit (ICU) admission, and requirement for mechanical ventilation, are heightened among patients who suffer a secondary bacterial infection in the aftermath of influenza (2, 3). Secondary bacterial infection requiring antibiotic treatment can in turn amplify the risk of further infections through the disruption of gut microbiota (4, 5), posing a serious concern in the era of rising antimicrobial resistance. Therapeutic approaches focusing on strengthening host defense against severe IAV infection could offer alternative treatment options. However, poor understanding of the immunological mechanisms governing severe influenza and the transition to post-viral bacterial infection hampers such efforts. Among mechanisms including epithelial cell death, influx of inflammatory cells, and impaired mechanical clearance, virus-induced depletion of the TR-AM pool has been suggested as a decisive step towards the establishment of post-influenza bacterial pneumonia (6, 7). TR-AM numbers remain relatively unchanged during homeostasis, with the main function of the cells being surfactant clearance and colonization (8). This tolerogenic programming, however, can be overridden by abundant viral presence, leading to a pro-inflammatory phenotypic switch, including extensive cytokine release and phagocytosis of viral particles and apoptotic cells (8-10). TR-AM loss, often observed after severe infection and notoriously known as the ‘TR-AM disappearance reaction’ (11, 12), characterizes IAV-induced pneumonia, aggravating lung injury, impairing host defense, and dramatically increasing IAV-associated mortality (13, 14). The specific pathomechanisms behind it remain, however, elusive.

The TNFSF involves a variety of structurally homologous ligands with multiple functions during development, homeostasis, and tissue response to injury (15). TNFSF14 or LIGHT [homologous to lymphotoxins, exhibits inducible expression and competes with Herpes Simplex Virus (HSV) glycoprotein D for herpes virus entry mediator (HVEM), a receptor expressed by T-lymphocytes] is widely expressed on cells of the hematopoietic compartment (16, 17). In the lung, TNFSF14 has been associated with airway remodeling in asthma, idiopathic pulmonary fibrosis, systemic sclerosis models (18, 19), and more recently, with disease severity in covid-19 (20, 21). Still, little is known regarding the pathomechanistic role of TNFSF14 in virus-induced pneumonia.

TNFSF14 binds to three different receptors; type I transmembrane lymphotoxin beta receptor (LTβR), HVEM, also known as TNF receptor superfamily 14 (TNFRSF14), and decoy receptor 3 (DcR3), which is only found in primates (15, 22). TNFSF14 receptors have a broad distribution on immune, stromal, and parenchymal cells (23), and orchestrate distinct intracellular pathways. The outcome of TNFSF14 crosslinking to its receptors, therefore, heavily relies on disease context and microenvironmental cues (15). Here, we sought to investigate the molecular mechanisms of post-influenza TR-AM loss and its consequences for host defense in a model of IAV and IAV/Spn infection, aiming at identifying any putative targets for immune-based pneumonia treatment options.

## Results

### Severe IAV infection increases susceptibility to secondary pneumococcal infection

To elucidate the pathomechanisms behind TR-AM death after severe IAV infection and its effect on the establishment of secondary bacterial pneumonia, we established a robust co-infection model. Disease severity after viral, bacterial, or co-infection was shown to vary in a pathogen- and infection dose-dependent manner. Orotratracheal (o.t.) infection of C57BL/6 wild-type (wt) mice with 500 foci-forming units (ffu) IAV A/PR/8/34 decreased mouse survival by 50% 14 days after infection, whereas intranasal infection with 2000 colony-forming units (cfu) Spn Serotype 3 (PN36 NCTC7978) neither affected survival (Figure 1A) nor led to any weight loss (Figure 1B). However, preexisting IAV infection seven days prior to pneumococcal infection caused a massive inflammatory response (Figure 1C) and a 100% lethal outcome (Figure 1A). Upon the use of lower IAV and Spn doses (250ffu/20cfu on day 7 post-IAV infection, pi), average survival was calculated at 37.5% (Figure 1A). Despite the low infection doses (250ffu IAV/20cfu Spn), bacterial load in the BALF of previously IAV-infected mice was remarkably high 48h after pneumococcal infection, whereas PBS-pretreated mice completely cleared pneumococci (Figure 1D), suggesting that prior IAV infection was associated with an impaired immune response against invading Spn. We therefore proceeded to characterize the leukocyte population landscape of the infected lung, as distinct immune cell populations and their interactions can differentially affect post-influenza bacterial clearance (7, 24, 25). Flow cytometric profiling of leukocyte composition in the BALF of IAV-infected mice (gating strategy depicted in Supplemental Figure 1A) revealed cell-specific kinetics in the alveoli over the course of IAV infection (Figure 1E). Of note, TR-AM numbers started declining on day 3 pi, reaching minimum levels on days 7-11 pi (Figure 1F), and depicting a drastic depletion of the pool. Similar results were shown for lung-tissue leukocyte populations (gating strategy in Supplemental Figure 1B), which could not be acquired through BAL due to their sessile nature (Supplemental Figure 2A) (26) or their extra-alveolar location (Supplemental Figure 2B-M). Impaired Spn clearance despite the abundance of professional phagocytes such as bone marrow-derived macrophages (BMDM) or neutrophils in the absence of TR-AM, which additionally showed a superior phagocytosis capacity on day 7 pi compared to BMDM (Fig. 1G), hinted at TR-AM importance for post-influenza transition to secondary bacterial pneumonia (7, 27). To address this, we sought to identify the molecular underpinnings of post-influenza TR-AM death.

**Figure 1.**
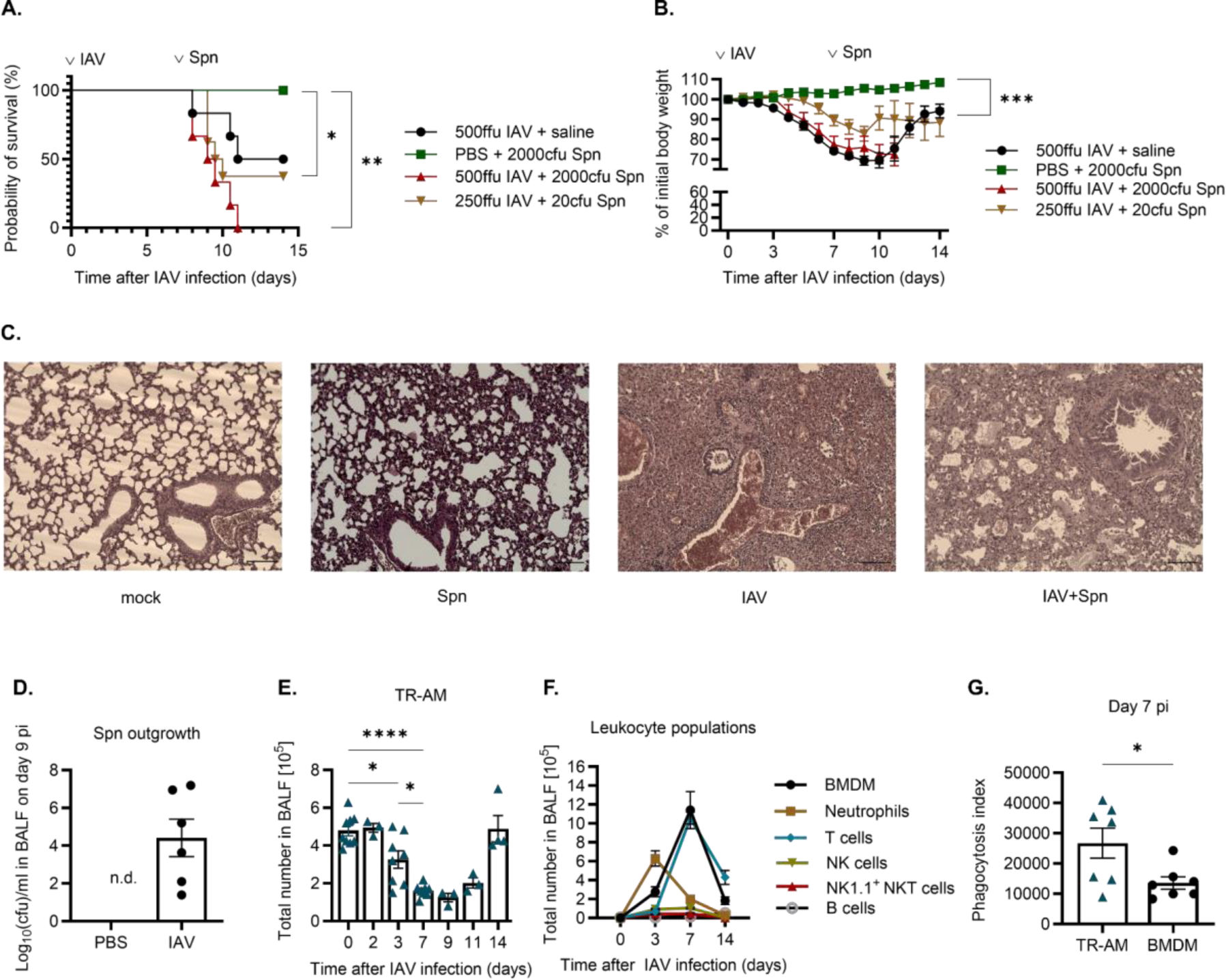
IAV infection increases susceptibility to secondary pneumococcal infection. **(A-B)** Survival (Figure 1A) and weight loss (Figure 2B) after infection of wt mice with IAV (n=6-8), followed by infection with Spn 7 days later. **(C)** Representative histological images of mock-infected mice (far left), mice infected only with Spn (left), mice infected only with IAV (right), or mice infected with IAV 7 days prior to Spn infection (far right), lungs harvested three days after Spn infection. Scale bar set at 100µm. **(D)** Bacterial load in the BALF of IAV-infected or PBS-treated mice nine days after infection with 250ffu IAV and 48h after infection with 20cfu Spn (mean ± SEM, n=6). **(E)** TR-AM population during the IAV infection course (n=3-9). **(F)** Leukocyte populations including neutrophils (n=7-9), BMDM (n=8-10), NK cells (n=6-11), T cells (n=3-10), NK1.1^+^ NKT cells (n=3-12), and B cells (n=5-9) in the BALF of infected mice on days 0,3,7, and 14 pi. Data shown are pooled from five different experiments, mean ± SEM is depicted. **(G)** Overall phagocytosis capacity of TR-AM compared to BMDM depicted as phagocytosis index, calculated by MFI FITC x % FITC+ cells, within each population on day 7 pi (means ± SEM, n=7). Significance was determined by log-rank (Mantel-Cox) test, unpaired t-test, and by 1-way ANOVA with Tukey’s posthoc test; *p<0.05, **p<0.01, ***p <0.001, ****p <0.0001.

### Post-influenza TR-AM death involves the activation of caspase-8

Direct viral infection can lead to epithelial cell apoptosis in IAV-induced pneumonia (28-30), posing the question whether this is also the driving mechanism of post-IAV TR-AM depletion. Flow cytometry analysis revealed only a small number of TR-AM being IAV-infected (quantified by virus hemagglutinin (HA) expression) with no significant increase over the course of infection (Figure 2A). The majority of HA-negative cells had been depleted by day 7 pi (Figure 2A), implying the involvement of a different mechanism with a much higher impact on TR-AM death. In accordance with that, when naïve TR-AM were *ex vivo* treated with virus- and cell-free BALF from IAV-infected wt mice from 7 pi (day of maximum depletion), we observed a significant decrease in TR-AM survival (Figure 2B) and an increase in caspase-3/7 (Figure 2C) and caspase-8 activity 24h after treatment (Figure 2D). Transcriptome analysis of flow-sorted HA-negative TR-AM on day 3 pi and on day 7 pi based on a cell-death gene array revealed an upregulation of multiple apoptosis-related genes, such as *Bax*, *Cd40lg*, and *Cflar*, and necrosis-related genes, including *Bmf*, *Commd4*, *Defb1*, and *Parp1*, compared to TR-AM from mock-infected mice (Figure 2E). Concomitant flow cytometry analysis revealed an increase in the percentage of apoptotic TR-AM over the infection course (Figure 2F, gating strategy in Supplemental Figure 3A). As a result, we raised the question whether post-influenza TR-AM survival could be improved by apoptosis inhibition. Following pre-incubation with 50µM of a specific caspase-3 (Z-DEVD-FMK) or caspase-8 inhibitor (Z-IETD-FMK), naïve TR-AM were treated with UV-inactivated BALF from infected mice. Whereas caspase-3 inhibition only showed a negligible protective effect, TR-AM death was completely abrogated in the caspase-8 inhibition group (Figure 2G), leading us to focus on caspase-8 for future experiments. Caspase-8 inhibition fully protected the TR-AM pool (Figure 2H), without affecting viral titers (Figure 2I). When mice were treated with daily subcutaneous (s.c.) injections of the caspase-8 inhibitor (schematic of experimental layout Figure 2J), an attenuated weight loss was observed up to 7 pi, compared to the DMSO group (Figure 2K). Caspase-8 inhibition further led to a significant reduction of TR-AM loss on day 7 pi (Figure 2L). Caspase-8 is a known orchestrator of cell death, which typically gets activated upon the crosslinking of a soluble ligand to a death receptor (31-33), which, together with the primarily virus-independent post-influenza TR-AM apoptosis we observed, hinted at a soluble ligand as a driver of TR-AM death early after IAV infection.

**Figure 2.**
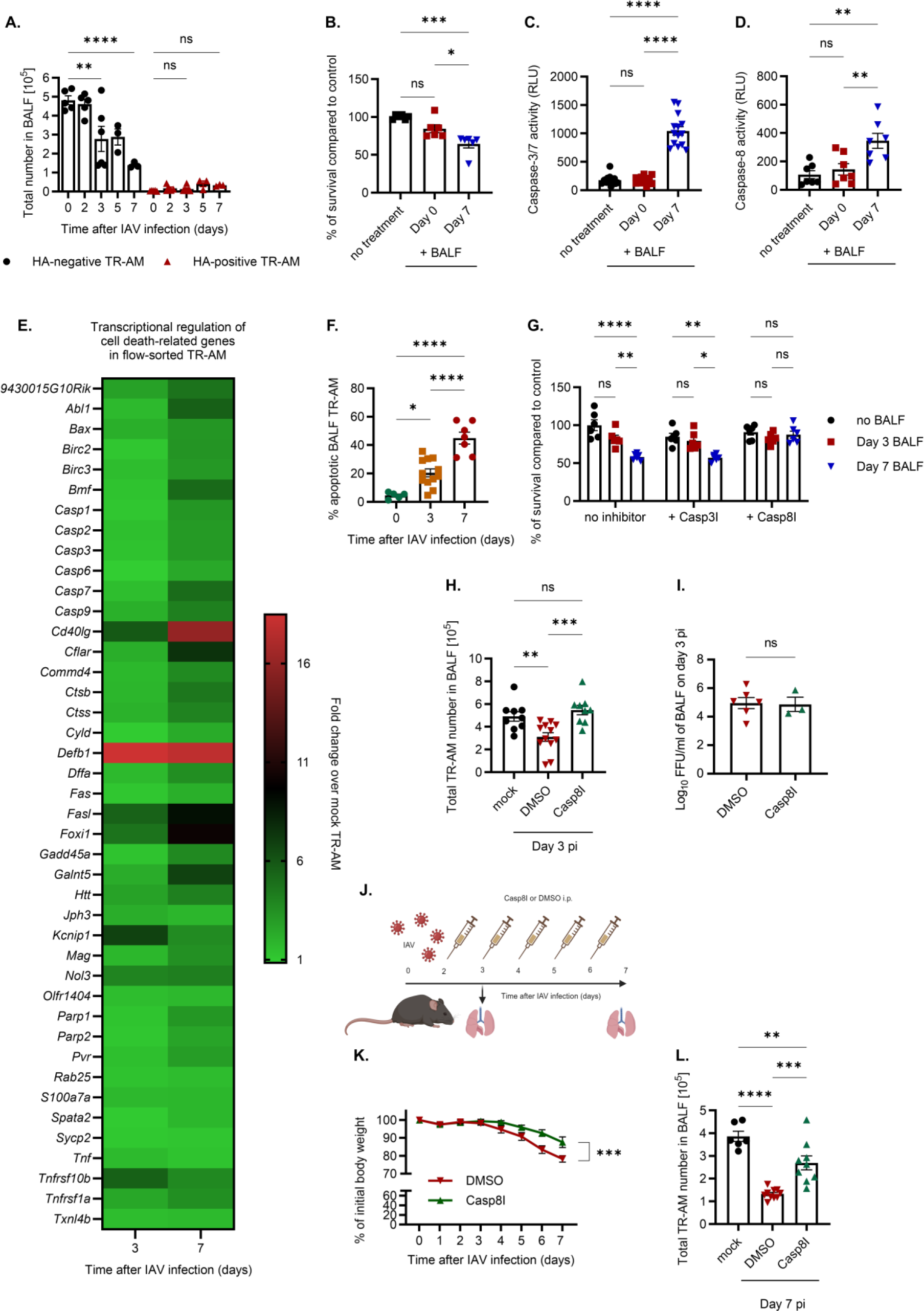
Caspase-8 is involved in virus-independent post-influenza TR-AM death. **(A)**Quantification of viral HA-negative and HA-positive TR-AM after IAV infection, mean ± SEM is depicted. **(B)** Survival of TR-AM following 24h of treatment with UV-irradiated BALF. Graphs represent means ± SEM, n=6. **(C-D)** Caspase-3/7 **(C)** and caspase-8 activity **(D)** following TR-AM treatment with BALF of mock- or IAV-infected mice. Caspase activity was measured as luminescence emission (means ± SEM of 6 independent experiments). **(E)** Heat map depicting average fold change of cell death-related genes. Transcriptomic analysis was performed in flow-sorted, HA-negative mock-infected and TR-AM collected on days 3 and 7 pi. **(F)** Percentage of apoptotic BALF on days 0, 3, and 7 pi (means ± SEM, n=5-12). **(G)** Colorimetric viability assay following treatment of naïve TR-AM with BALF from IAV-infected animals and pre-treatment with a specific caspase inhibitor (means ± SEM, n=6). **(H)** BALF TR-AM on day 3 pi following IAV infection and caspase-8 inhibition (means ± SEM, n=9-12). **(I)** Viral titers in the BALF of infected mice on day 3 pi, following IAV infection and caspase-8 inhibition (means ± SEM, n=3-6). **(J)** Experimental layout for caspase-8 inhibition in vivo experiments. **(K)** Weight loss after IAV infection and caspase-8 inhibition, n=8-11. **(L)** BALF TR-AM on day 7 pi (means ± SEM, n=7-10). Significance was determined by t-test of the Areas Under Curve (AUC) for the DMSO and Casp8I group in **(K)**, 1-way, or 2 -way ANOVA with Tukey’s posthoc test; *p<0.05, **p<0.01, ***p <0.001, ****p <0.0001.

### IAV pneumonia sensitizes TR-AM to TNFSF14 ligation

Death-inducing members of the TNFSF have been associated with promoting alveolar epithelial cell death and driving post-IAV lung injury in a series of studies (24, 34, 35). As such, we hypothesized that a TNFSF member could be involved in post-IAV TR-AM death and analyzed gene expression patterns of receptors and ligands belonging to the TNFSF signaling network in flow-sorted, HA-negative TR-AM from mock-infected and infected mice on day 3 and day 7 pi. *Tnfrsf14*, a receptor for *Tnfsf14*, showed a significant upregulation at both time points (Figure 3A, fold change of 8.59 over mock, p 0.049, on day 3, and 305.13, p 0.009, on day 7 pi). TNFRSF14 demonstrated a significant increase in mRNA and TR-AM cell surface protein expression levels over the infection course or after *ex vivo* stimulation of TR-AM with BALF from IAV-infected mice (Figure B-D). No significant transcriptional changes could be observed for *Ltβr*, the competitor receptor for TNFSF14, though protein expression on TR-AM also increased on day 7 pi (Figure 3B-C, E). Regarding TNFSF ligand expression, we confirmed previous reports on IAV-induced upregulation of *Tnfsf10* (fold change of 11.35 over mock, p 0.036, on day 3 pi, and 29.61, p 0.030, on day 7 pi) in lung macrophages (34, 35), whereas *Tnfsf15* and *Tnfsf14* were moderately increased (*Tnfsf15*: fold change of 7.21 over mock, p 0.140, on day 3 pi, and 11.18, p 0.004, on day 7 pi, *Tnfsf14*: fold change of 5.36 over mock, p 0.039, on day 3 pi, and 9.44, p 0.009, on day 7 pi) (Figure 3F). Overall, severe IAV infection was linked to a total increase in TNFSF14 in the mouse lung, as shown by qPCR (Figure 3G), IHC (Figure 3H), and for soluble TNFSF14 by ELISA in the BALF of infected animals (Figure 3I). In accordance with that, we observed a significant increase in soluble TNFSF14 in the BALF of patients with influenza or COVID-19 ARDS, compared to control patients who underwent routine bronchoscopy for diagnostic purposes and revealed normal BALF cellularity (Figure 3J). Based on the distinct kinetics of the two TNFSF14 receptors on TR-AM after IAV infection, we aimed at dissecting any differential roles in post-influenza TR-AM fate. To this end, we treated naïve wt, *tnfrsf14^-/-^*, and *ltβr^-/-^* TR-AM with virus-free BALF for 24h. Caspase 3/7 activity was significantly lower in the *ltβr^-/-^* group (Figure 3K), indicating a potential protective effect against soluble death-inducing ligands in the absence of the receptor. In accordance with that, IAV-infected *ltβr^-/-^* mice demonstrated a less dramatic drop in TR-AM numbers, compared to wt and *tnfrsf14^-/-^* mice (Figure 3L) as well as an overall attenuated weight loss on days 7 and 8 pi (Figure 3M). The distinct effects of the two receptors on TR-AM survival and the upregulation of the common TNFSF14 ligand in IAV-infected lungs led us to examine TNFSF14 closer in a further experimental series.

**Figure 3.**
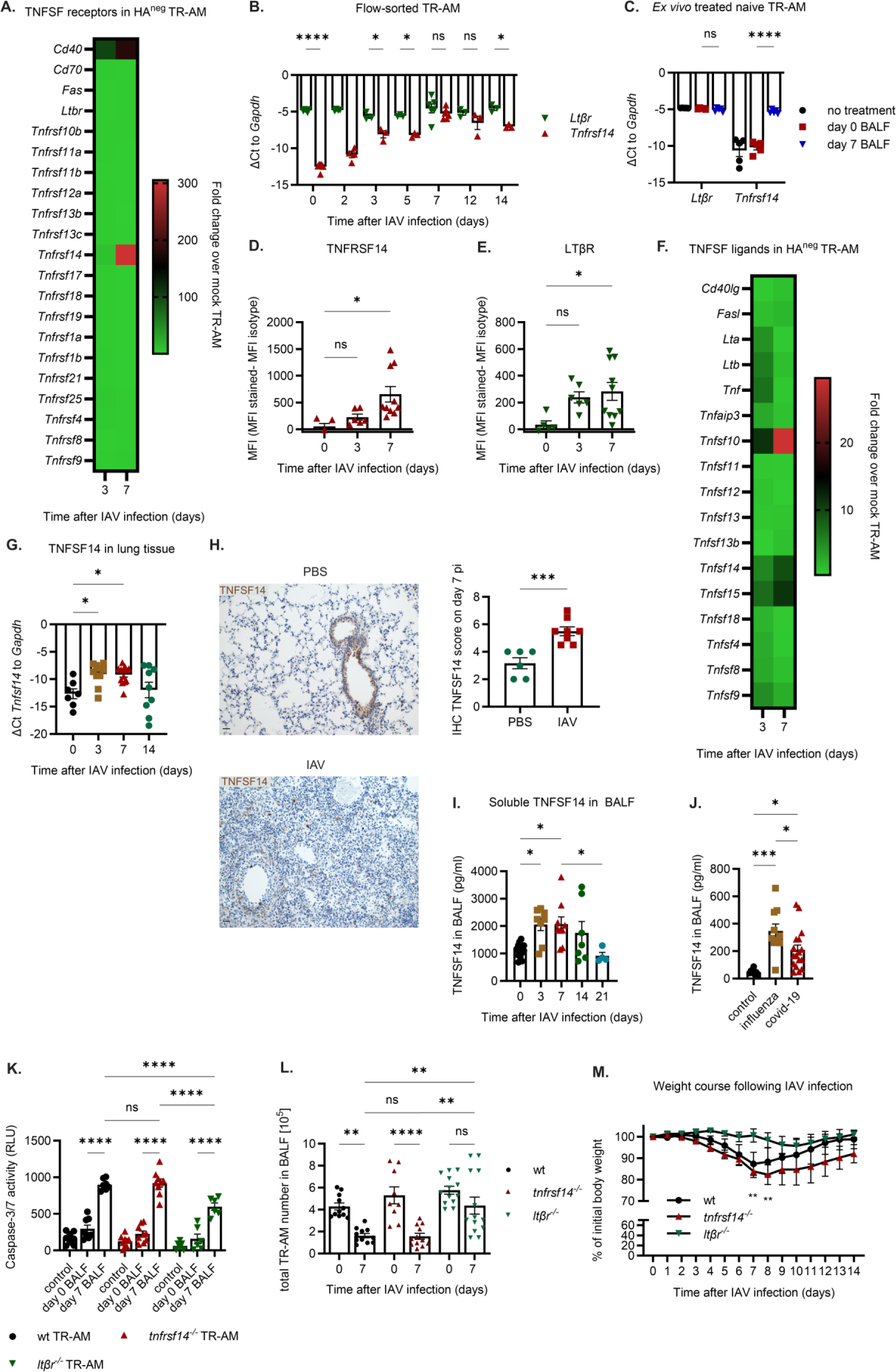
IAV infection leads to an increased expression of the TNFSF14 ligand-receptor axis in the lung. **(A)**Fold change of receptor genes belonging to the TNF superfamily (TNFSF) in BALF TR-AM on days 3 and 7 pi, over mock TR-AM (n=5-6). **(B)** *Tnfrsf14* and *Ltβr* gene expression in flow-sorted TR-AM after IAV infection. Graph represents means ± SEM, n=3-6. **(C)** *Tnfrsf14* and *Ltβr* gene expression in naïve TR-AM 24h after treatment with BALF from IAV-infected mice (means ± SEM, n=5 per condition). **(D-E)** Flow cytometry analysis for TNFRSF14 **(D)** and LTβR **(E)** expression on TR-AM during the early IAV infection course. MFI normalized to isotype-stained samples (means ± SEM, n=5-10). **(F)** Fold change of TNFSF ligand genes in BALF TR-AM on days 3 and 7 pi over mock TR-AM, n=5-6. **(G)** *Tnfsf14* gene expression in the lungs of IAV-infected animals (means ± SEM, n=7-8). **(H)** Lung section IHC analysis for TNFSF14 expression after mock (PBS) or IAV infection. Scale bar set at 25μm (means ± SEM, n=6-8). **(I)** Soluble TNFSF14 in the BALF of IAV-infected mice measured by ELISA (means ± SEM, n=3-8). **(J)** Soluble TNFSF14 in the BALF of patients with severe viral pneumonia and controls (means ± SEM, n=8-17). **(K)** Caspase-3/7 activity after 24h incubation of naïve wt, *tnfrsf14^-/-^*, and *ltβr^-/-^* TR-AM with BALF from IAV-infected mice. Graph represents means ± SEM, n=3-8. **(L)** BALF TR-AM numbers of wt, *tnfrsf14^-/-^* and *ltβr^-/-^* mice on day 0 and day 7 pi (means ± SEM, n=9-13). **(M)** Body weight of wt, *tnfsf14^-/-^*, and *ltβr^-/-^* mice over the course of IAV infection up to day 7 pi(means ± SEM of weight at each time point, n=5-6). Significance was determined by unpaired t-test for IHC score comparison, 1-way ANOVA with Tukey’s posthoc test, and Kruskal-Wallis test followed by Dunn’s post hoc comparison test for **(J)**; *p<0.05, **p<0.01, ***p <0.001, ****p <0.0001.

### TNFSF14 drives the depletion of the TR-AM pool in IAV-induced pneumonia

To test whether TNFSF14 could directly induce TR-AM death, we treated naïve murine (Figure 4A) and human BALF TR-AM (Figure 4B) with different concentrations of recombinant TNFSF14 (rTNFSF14) for 24h. Treatment with 500ng/ml rTNFSF14 led to an average of 25-35% decrease in TR-AM survival compared to the control PBS/BSA group, as measured by a cytotoxicity assay (Figure 4A-B), and an increase in caspase-3/7 activity, which was even more pronounced when the same setup was performed with TR-AM isolated on day 3 post IAV infection (Figure 4C). In addition to our *in vitro* experiments, we performed o.t. application of rTNFSF14 to mice infected with 500ffu and analyzed TR-AM numbers and apoptosis rates on day 3 pi (schematics in Figure 4D). We observed a remarkable increase in the number of Annexin V^+^ TR-AM in the BALF (Figure 4E) as well as a significant aggravation of the already diminished BALF and lavaged lung-tissue TR-AM numbers (Figure 4F-G), compared to the PBS-treated, IAV-infected control, proving the deleterious role of TNFSF14 for the TR-AM population. No further differences could be detected in the numbers of other leukocyte populations, depending on rTNFSF14 treatment (Supplemental Figures 4A-E). Concomitantly, TR-AM loss was completely abrogated in the BALF and in the lavaged lung tissue of *tnfsf14^-/-^* mice, as opposed to wt mice (Figures 5A-B). This result could not be reproduced in *tnfsf10^-/-^* mice (Supplemental Figure 5A), despite TNFSF10 being one of the pro-apoptotic ligands highly expressed in the BALF during IAV infection (34, 35), suggesting that TR-AM apoptosis induction was ligand-specific. Flow-sorted BALF TR-AM from *tnfsf14^-/-^* mice showed lower fold changes of apoptosis-(Figure 5C-D), necrosis (Supplemental Figures 5B-C), and autophagy-related genes (Supplemental Figures 5D-E) on day 3 and day 7 pi over non-infected controls, as determined per gene array analysis for cell-death related genes, compared to wt mice. Unlike wt BALF treatment, caspase-3/7 activity was not increased upon TR-AM treatment with BALF from *tnfsf14^-/-^* mice (Figure 5E). No difference could be detected in viral titers on day 3 pi (peak of viral replication in this model (36), Supplemental Figure 5F) or in the amount of EpCAM+ lung epithelial cells (suggesting no induction of epithelial cell apoptosis by TNFSF14, Supplemental Figure 5G). There was also no evidence of TNFSF14-dependent cell death of further BALF leukocyte populations (Supplemental Figure 5H-L), suggesting that it was confined to the TR-AM compartment. Alongside, absence of the ligand resulted in decreased weight loss, hinting at a beneficial effect of TNFSF14 blockade (Figure 5F). Aiming at a therapeutic approach, we tested outcome after systemic treatment with a neutralizing anti-TNFSF14 (clone 3D11) antibody. We observed higher BALF and lung tissue TR-AM numbers on day 7 pi (Figures 5G-H) and an attenuated weight loss in the anti-TNFSF14 antibody group (Figure 5I) and confirmed lower caspase-3 activity in TR-AM after *ex vivo* wt BALF treatment (Figure 5J), highlighting TNFSF14 as the driver of post-influenza TR-AM death.

**Figure 4.**
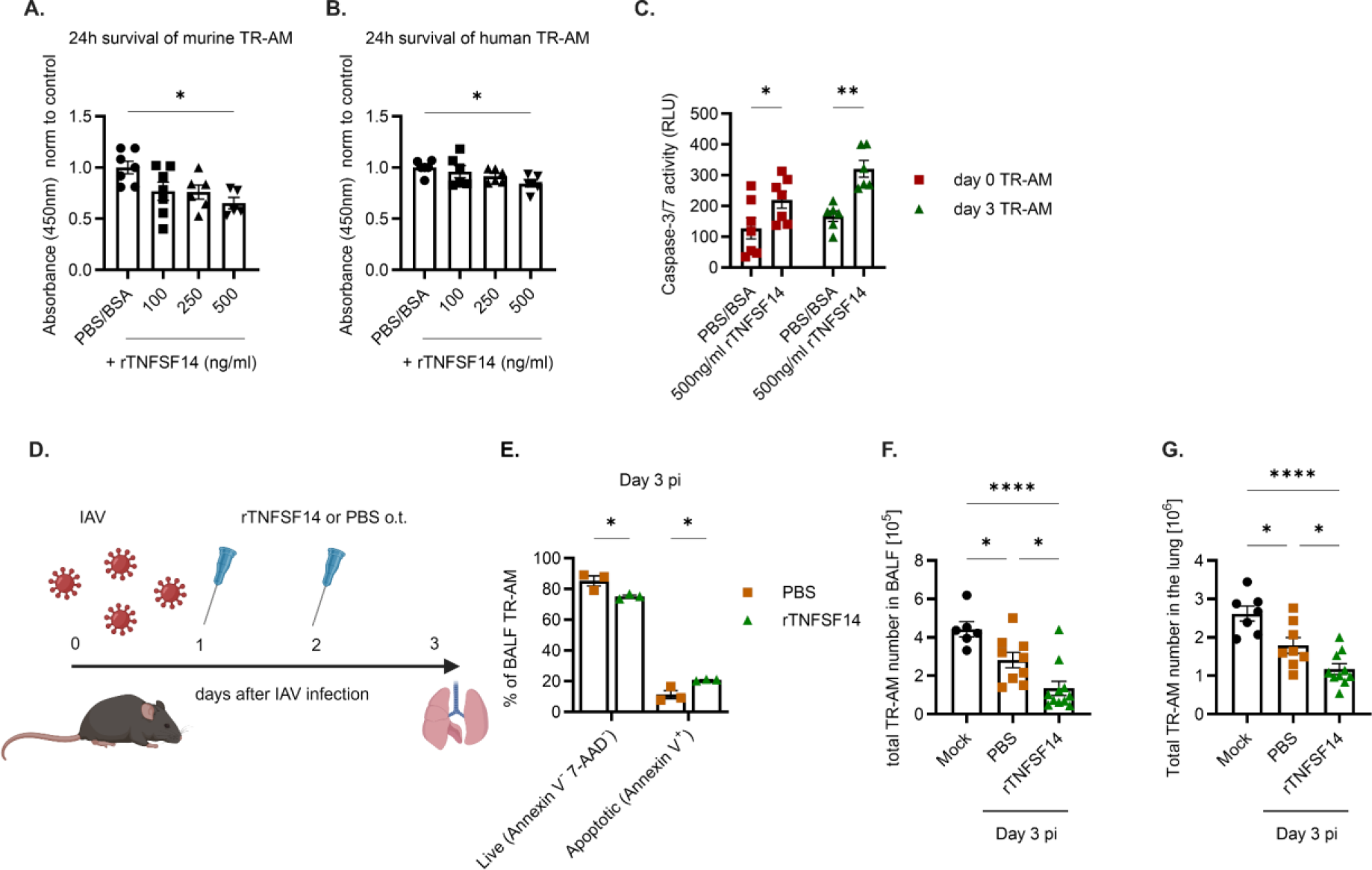
TNFSF14 treatment aggravates post-influenza TR-AM loss. **(A-B)** Colorimetric assay depicting cell survival after 24h treatment of murine **(A)** and human **(B)** TR-AM with different doses of recombinant TNFSF14 (rTNFSF14) or a PBS/BSA control. Graph represents means ± SEM, n=5-7. **(C)** Caspase-3/7 activity after 24h treatment of BALF TR-AM with 500ng/ml rTNFSF14 (means ± SEM, n=6-7). **(D)** Schematics of rTNFSF14 application in IAV-infected mice with analysis performed on day 3 pi. **(E)** Percentage of live (Annexin V^-^ 7-AAD^-^) and apoptotic (Annexin V^+^) TR-AM (n=3) on day 3 pi after rTNFSF14 treatment. **(E-F)** Total BALF (n=6-11) **(E)** and lung-tissue TR-AM numbers **(F)** on day 3 pi after rTNFSF14 treatment (mean ± SEM, n=7-10). Significance was determined by 1-way or 2-way ANOVA with Tukey’s posthoc test; *p<0.05, **p<0.01, ***p <0.001, ****p <0.0001.

**Figure 5.**
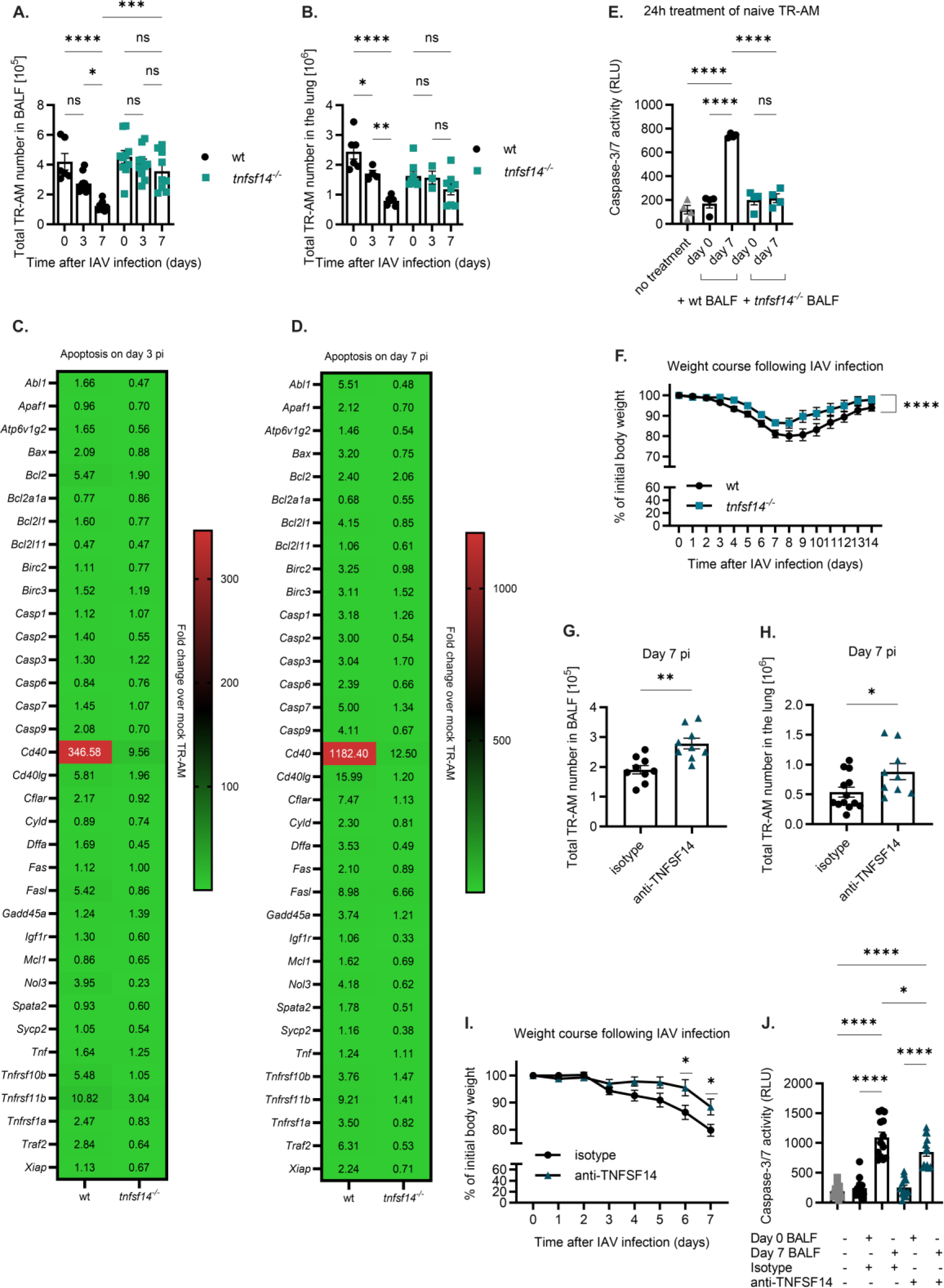
Post-influenza TR-AM loss can be prevented through directed targeting of the TNFSF14 ligand. **BALF** TR-AM **(A)** and sessile TR-AM **(B)** in wt and *tnfsf14^-/-^* mice after IAV infection (means ± SEM, n=8-12). **(C-D)** Heat maps depicting fold change of apoptosis-related genes on day 3 pi **(C)** and day 7 pi **(D)** over mock-infected wt and *tnfsf14^-/-^* TR-AM, n=3-5, as determined per gene array analysis. **(E)** Caspase-3/7 activity after TR-AM treatment with wt and *tnfsf14^-/-^* BALF. Graph represents means ± SEM, n=4 per condition. **(F)** Body weight of wt and *tnfsf14^-/-^* mice over the course of IAV infection. Graph represents means ± SEM of weight at each time point, n=12-13. **(G-H)** BALF **(G)** and lung tissue **(H)** TR-AM on day 7 pi after treatment with an anti-TNFSF14 antibody or isotype control on day 2 pi. Graph represents means ± SEM, n=9-13. **(I)** Body weight of anti-TNFSF14 and isotype-treated wt mice up to day 7 after IAV infection. Graph represents means ± SEM of weight at each time point, n=11-12. **(J)** Caspase-3/7 activity after incubation of naïve TR-AM with BALF pre-treated with an anti-TNFSF14 antibody. Data pooled from n=4 experiments (means ± SEM, n=13). Significance was determined by unpaired t-test, 1-way, or 2-way ANOVA with Tukey’s posthoc test; *p<0.05, **p<0.01, ***p <0.001, ****p <0.0001.

### TNFSF14 is released by neutrophils during the acute IAV infection phase

To identify the cellular source of TNFSF14 during IAV pneumonia, we analyzed expression of the transmembrane form of TNFSF14 on leukocytes (CD45^+^ cells), epithelial cells (EpCAM^+^ cells), mesenchymal cells (MC), and endothelial cells (CD31^+^) in the lungs of mock- and IAV-infected mice on day 7 pi. TNFSF14 expression was significantly increased in epithelial cells and leukocytes (Figure 6A). This was in accordance with our IHC data, which revealed a prominent signal increase within the leukocyte-infiltrated interstitium and alveolar space on day 7 pi (Figure 6B). Soluble TNFSF14 was detected in the serum of infected mice as early as day 3 pi, further suggesting that blood-derived immune cells contributed to the increase in TNFSF14 levels within the lung (Figure 6C). Combined single cell (sc)RNA-Seq analysis of pre-sorted CD45+ cells from whole lung digests on days 3 and 7 pi revealed a total of 14 immune cell clusters on day 3 pi, including TR-AM, BMDM, interstitial macrophages (IM), B and T cells, conventional dendritic cells (cDCs), plasmacytoid DCs (pDCs), monocyte-derived DCs (moDCs), NK cells, three monocyte clusters with different gene signatures: monocytes 1 (*Ccr2, Ly6a2, F13a1, Mgst1, Aldh2*), monocytes 2 (*Tgfb1, Sirpb1c, Otulin1, Plcg2, Zfp710*), and monocytes 3 (*Eno3, Cd300e, Agpat4, Rbpms, Slc12a2*), and two neutrophil clusters: neutrophils 1 (*Ier5, Ier3, Smox, Gm8995, Ccrl2*) and neutrophils 2 (*Picalm, Jund, Hmgb2, Map1lc3b, Slc2a3*). Eleven clusters were identified on day 7 pi, including TR-AM, BMDM, IM, B cells, CD4+, CD8+ and proliferating T cells, pDCs, cDCs, NK cells, and neutrophils (Figure 6D-E). On both time points, neutrophils were revealed as the main leukocyte population expressing *Tnfsf14*, with a minor contribution from NK and T cells. *Ltβr* gene expression was higher than *Tnfrsf14* in all monocyte/macrophage populations, including TR-AM (Figure 6D-E). qPCR analysis further revealed an upregulation of *Tnfsf14* in neutrophils isolated from the peripheral blood at day 2 pi, compared to non-infected mice (Figure 6F), confirmed in BALF neutrophils of patients with IAV ARDS (Figure 6G). We then tested whether neutrophil depletion via i.p. administration of an anti-Ly6G antibody (37) would improve post-influenza TR-AM survival. Following successful neutrophil depletion (Figure 5H and Supplemental Figures 6A-C), mice demonstrated an attenuated weight loss up to day 7 pi (Figure 6I). Neutrophil-depleted mice presented lower *tnfsf14* mRNA (Figure 6J) and soluble protein levels (Figure 6K), compared to the isotype-treated group, with higher BALF and lung-tissue TR-AM numbers on day 7 pi (Figure 6L-M). No significant differences in further BALF immune cell populations (Supplemental Figures 6 D-H), demonstrating the depletion specificity of the anti-Ly6G antibody.

**Figure 6.**
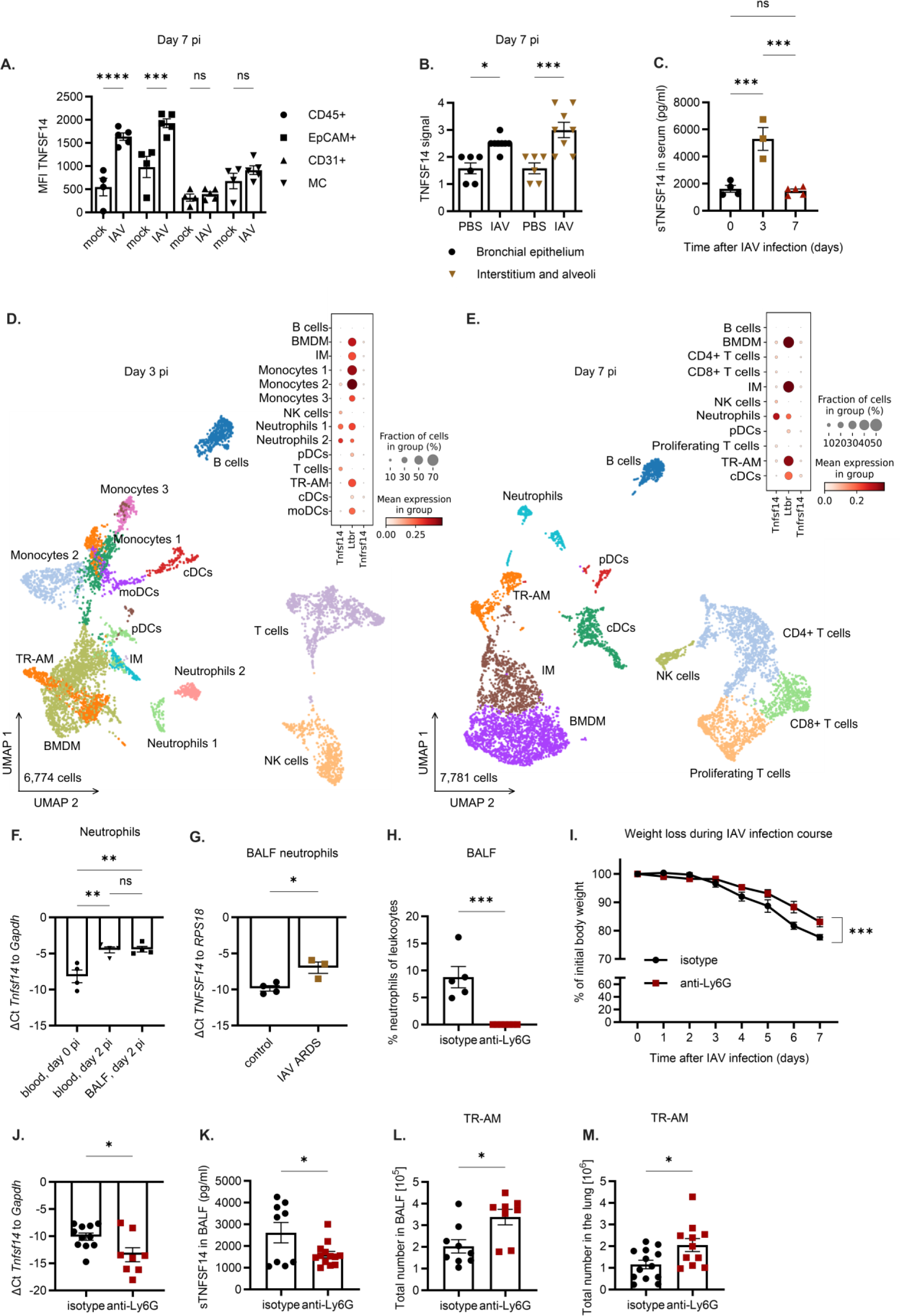
Neutrophils are the main cellular source of TNFSF14 during IAV infection. **(A)**MFI for TNFSF14 expression in leukocytes (CD45^+^ cells), epithelial cells (EpCAM^+^ cells), endothelial cells (CD31^+^ cells), and mesenchymal cells (MC) in the lungs of non-infected and infected mice on day 7 pi, normalized to the MFI of an isotype control. Graph represents means ± SEM, n=4-5. **(B)** IHC quantification of TNFSF14 signal in different lung regions of mock- and IAV-infected wt mice on day 7 pi (means ± SEM,n=3-4). Scale bar set at 25µm. **(C)** Soluble TNFSF14 in the serum of IAV-infected mice on days 0,3, and 7 pi (means ± SEM, n=3-5). **(D-E)** ScRNA-Seq analysis of flow-sorted leukocytes on day 3 **(D)** and day 7 pi **(E)** after IAV infection. Monocyte and neutrophil clusters on day 3 pi were characterized according to the gene signature of the top 5 uniquely expressed genes in each cluster. Dot plots depict *Tnfsf14, Ltβr,* and *Tnfrsf14* expression among the different clusters. **(F)** qPCR analysis for *Tnfsf14* expression ín murine neutrophils isolated from the blood and the BALF of IAV-infected mice (means ± SEM, n=4). **(G)** qPCR analysis for *TNFSF14* expression in neutrophils isolated from the BALF of patients with severe IAV-induced ARDS compared to non-IAV controls (means ± SEM, n=3-4). **(H)** Neutrophils depicted as percentage of leukocytes in the BALF of IAV-infected animals after the i.p. administration of 200μg/200µl PBS of a neutrophil-depleting antibody (anti-Ly6G, clone 1A8) or an isotype control (clone 2A3). Graphs represent means ± SEM, n=5-8. **(I)** Body weight as percentage of initial weight after IAV infection and neutrophil depletion, n=7-9. **(J-K)** qPCR analysis, n=8-11 **(J)**, and ELISA, n=9-13 **(K)**, for TNFSF14 expression after neutrophil depletion on day 7 pi after IAV infection. Graph represents means ± SEM. **(L-M)** BALF **(L)** and lung **(M)** TR-AM of IAV-infected animals on day 7 pi after anti-Ly6G or isotype control treatment (means ± SEM, n=8-13). Significance was determined by unpaired t-test, 1-way, or 2-way ANOVA; *p<0.05, **p<0.01, ***p <0.001, ****p <0.0001.

### Outcome of post-influenza pneumococcal pneumonia is improved by targeting the TNFSF14 ligand-receptor axis

Having established that TR-AM loss was driven by the TNFSF14-LTβR axis in severe IAV infection, we tested whether maintenance of the TR-AM pool would improve outcome in the co-infection model (Figures 1A-D). Indeed, *tnfsf14^-/-^* mice showed significantly improved survival and reduced weight loss (Figures 7A-B) after secondary Spn infection, associated with reduced pneumococcal burden in BALF on day 9 pi (Figures 7C and D), whereas no difference could be shown for spleen bacterial burden (Supplemental Figure 7A). Despite a massive neutrophil influx in both groups (Supplemental Figure 7B), *tnfsf14*^-/-^ mice maintained their TR-AM numbers (Figure 7E). No significant differences could be observed in the phagocytosis capacity and the percentage of phagocytic TR-AM at the time point of pneumococcal infection (Supplemental Figures 7C-D). Based on the lack of difference in overall phagocytic capacity, we concluded that the improved bacterial clearance we observed in the *tnfsf14^-/-^* mice was mainly attributed to the higher number of TR-AM surviving the IAV infection. Conversely, the improved outcome of *tnfsf14^-/-^* mice could be completely abrogated when TR-AM were clodronate-depleted (Figure 7F). Treatment with the neutralizing anti-TNFSF14 antibody led to a similar improvement in survival and weight loss (Figures 7G-H), supporting the hypothesis that post-influenza maintenance of the TR-AM pool was key to survive secondary pneumococcal infection. Finally, we performed orthotopic transfer of wt, *tnfrsf14^-/-^*, or *ltβr^-/-^* TR-AM on day 3 pi in the co-infection model (schematics in Figure 7I). Pneumococcal superinfection was lethal in mice which had received no cells or *tnfrsf14^-/-^* TR-AM, whereas transfer of *ltβr^-/-^* TR-AM rescued 75% of mice (Figure 7J). Transfer of wt TR-AM only slightly increased survival, indicating that transferred wt TR-AM experienced TNFSF14-dependent apoptosis after transfer at day 3 pi. *Ltβr^-/-^* TR-AM additionally showed a superior phagocytosis capacity for Spn compared to wt TR-AM when infected *ex vivo* (Figure 7K), which may have further contributed to the improved survival observed in the *ltβr^-/-^* TR-AM recipient group (Figure 7J). Based on these results, we conclude that targeting the TNFSF14 signaling axis could revert TR-AM death in the aftermath of severe IAV-induced lung injury and thus prevent the transition to secondary bacterial pneumonia (Figure 7L).

**Figure 7.**
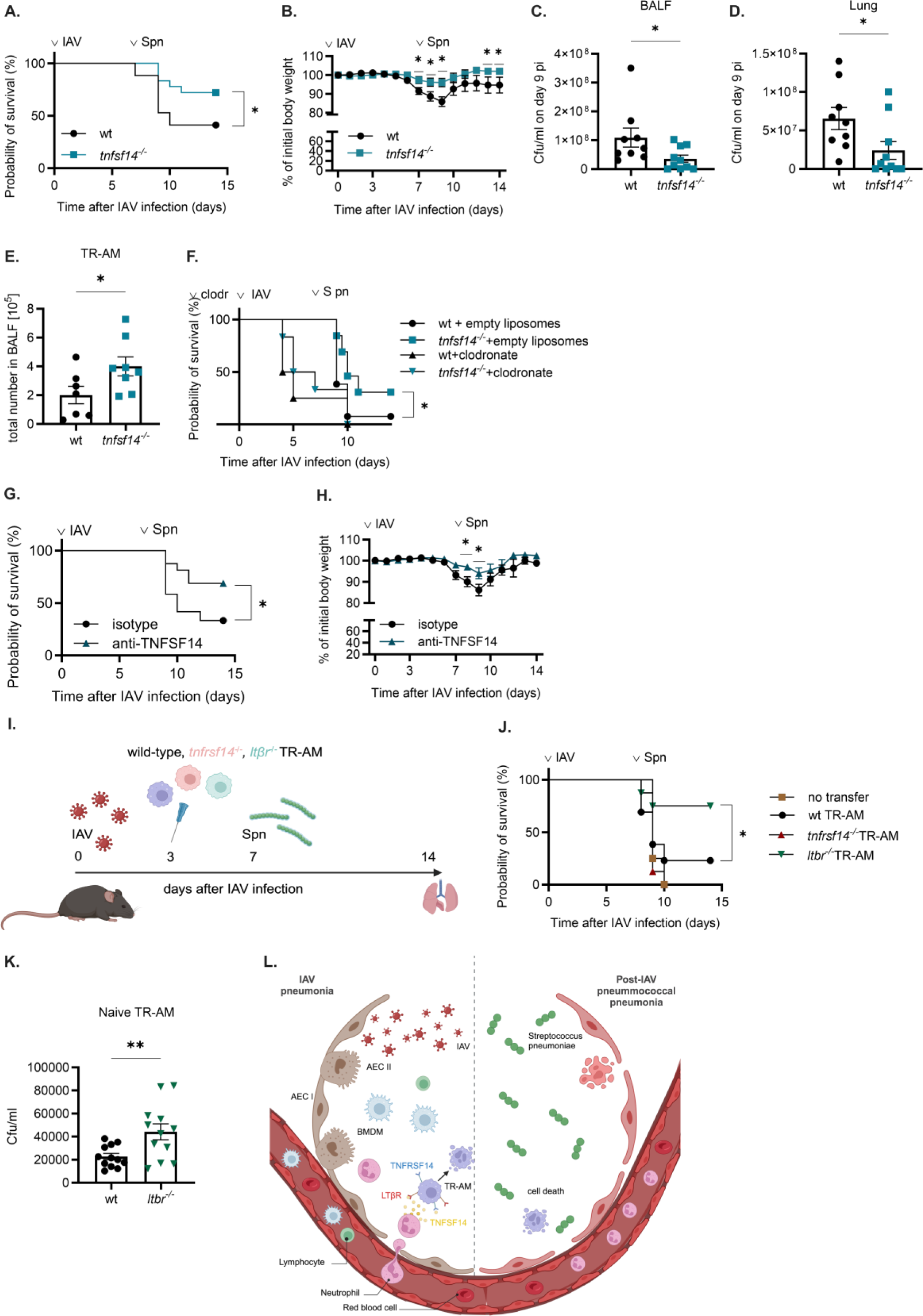
Severity of post-influenza pneumococcal pneumonia is attenuated in the absence of TNFSF14. (A-B) Survival **(A)** and weight loss **(B)** after infection of wt and *tnfsf14^-/-^* mice with 250ffu IAV and 20cfu Spn on day 7 post-IAV infection, (means ± SEM, n=17-18). **(C-D)** Bacterial burden in the BALF **(C)** and in the lungs **(D)** of wt and *tnfsf14^-/-^* mice 9 days after IAV infection and 48h after Spn infection (means ± SEM, n=7-9). **(E)** Total TR-AM numbers in the BALF of wt and *tnfsf14^-/-^* mice on day 9 pi after IAV infection and 48h after Spn (means ± SEM, n=7-8). **(F)** Survival of wt and *tnfsf14^-/-^* mice following co-infection and o.t. application of 70µl clodronate or empty liposomes two days prior to IAV infection (means ± SEM, n=4-13). **(G-H)** Survival **(G)** and weight loss **(H)** following IAV/Spn co-infection and i.p. administration of 500μg of a blocking anti-TNFSF14 antibody or an isotype control on day 2 pi (means ± SEM, n=12-16). **(I)** Schematics of experimental layout for IAV/Spn co-infection and adoptive transfer (a.t.) of naïve TR-AM on day 3 pi. **(J)** Survival of wt mice upon IAV/Spn co-infection and a.t. of wt, *tnfrsf14^-/-^*, or *ltβr^-/-^* TR-AM on day 3 pi (means ± SEM, n=4-12). **(K)** Bacterial load in lysed wt and *ltβr^-/-^* naïve TR-AM after *ex vivo* infection with Spn at an MOI 1000 (means ± SEM, n=12). Significance was determined by log-rank (Mantel-Cox) test, unpaired t-test, 1-way and 2-way ANOVA with Tukey’s posthoc test; *p<0.05, **p<0.01. **(L)** Proposed hypothesis: Severe IAV-induced pneumonia is characterized by massive leukocyte recruitment, including neutrophils. Once in the alveoli, neutrophils start releasing TNFSF14, which is sensed by TR-AM through ligation to surfaced-expressed receptors TNFRSF14 and LTβR, culminating in TR-AM death, which increases host susceptibility to post-influenza pneumococcal pneumonia.

## Discussion

Lower respiratory tract infections (LRTIs) remain one of the leading causes of death worldwide, with a severe toll on public health even prior to the pre-covid-19 era (38). IAV infection holds a particular role among LRTIs due to a variety of potential complications, with secondary bacterial infection being one of the most severe, as it dramatically increases the risk of respiratory failure, need for ICU treatment, and overall mortality (1, 28, 39). With no causative pharmacological treatment for pneumonia-related lung injury currently available, studies have been focusing on improving our understanding of the mechanisms which govern severe IAV pneumonia and favor the transition to post-influenza secondary infection. IAV-associated epithelial cell death allows disseminating bacteria to enter the inflamed tissue, where excessive inflammatory cell influx, exuberant cytokine release, and interferon-driven dampening of phagocyte antibacterial properties, further compromise host defense (6, 40, 41). Fibrin deposition, impaired mechanical clearance, and an abundance of newly available glycoconjugates additionally support bacterial outgrowth (42). Influenza-caused microbial dysbiosis in the respiratory and in the gastrointestinal tract changes inter-microbial dynamics, which may further lead to the selection of highly virulent bacterial species (43). With TR-AM safeguarding the alveoli as the first line of defense, IAV-induced TR-AM death can be viewed as one of the key events towards a compromised host defense. Patient studies utilizing scRNA-Seq and flow cytometry data have demonstrated that severe viral infections lead to the depletion of the TR-AM pool and the replenishment of the niche by BMDM (44, 45). Animal and *in silico* models of IAV pneumonia have positively correlated the depletion phase of TR-AM with the period of maximum susceptibility to secondary bacterial pathogens (7, 46), yet the mechanisms behind TR-AM depletion remain poorly understood. In this study, we identified a previously understudied member of the TNFSF as a driver of TR-AM depletion in the context of IAV pneumonia.

TR-AM numbers started declining as early as day 3 pi and intensified on day 7 pi, while other leukocyte populations gradually appeared in the alveoli as a response to the viral infection. TR-AM fate after acute infection is proposed to be dictated by cell death-inducing mechanisms, impaired self-renewal capacity, or loss of pro-survival signals from the neighboring epithelium due to epithelial destruction (11, 12). The extent of TR-AM depletion, the duration, and the intensity of the inflammatory response, determine the cellular composition and (re)programming of lung-resident cells in the aftermath of infection (47). Our own previous data indicates that partial TR-AM loss enables the recruitment of circulating BMDM, which are essential for post-viral repair through their transitioning into pro-homeostatic phenotypes (48). Co-existence of newly recruited BMDM and surviving original TR-AM is the outcome of a balanced immune response, which culminates in BMDM-orchestrated tissue repair and return to homeostasis, assisted by the tolerogenic functions of TR-AM, such as neutrophil efferocytosis, aimed at restricting epithelial damage (49, 50).

Infection severity determines the extent of TR-AM depletion, with a dramatic loss being observed upon severe IAV pneumonia, as demonstrated in our model. Recruitment of pro-inflammatory immune cells as a consequence of the disrupted gas-blood barrier and the abundance of chemokines present contribute to viral clearance, but can also signify the transition to a dysbalanced immune response (51). The highly pro-inflammatory programming of BMDM, for instance, can aggravate local injury and promote aberrant lung remodeling (24, 36), while uncontrolled neutrophil migration and dysregulated activation positively correlate with disease severity and negative patient outcome in severe influenza cases (52, 53). Regarding post-viral secondary infection, we observed that immediate recruitment of professional phagocytes such as BMDM and neutrophils was not sufficient to prevent bacterial dissemination in the aftermath of TR-AM depletion. This is in accordance with previously published data, which demonstrated a high susceptibility to secondary pneumococcal infection during the TR-AM depletion phase in a mouse co-infection model (7). The near-complete TR-AM loss in our model can thus be viewed as collateral damage within a microenvironment of abundant death-inducing signals, resulting in overshooting BMDM recruitment and creating a window of increased immunological vulnerability, where secondary pneumococcal infection proved lethal.

Through transcriptomics and flow cytometry analysis we identified apoptosis as the main pathway driving post-influenza TR-AM death, highly independent of direct viral infection, suggesting the involvement of a death-inducing ligand. Apoptosis supports viral propagation during early infection, as it promotes viral release and infection of new targets (30, 54). As a host defense mechanism, apoptosis can deter viral dissemination and promote viral clearance through the elimination of infected cells (55, 56). Leukocyte- and virus-driven apoptosis of alveolar epithelial cells, on the other hand, compromises the gas-blood barrier and impairs gas exchange (34, 36). Apoptosis inhibition through the use of specific caspase inhibitors can, therefore, affect the course of infection in a multitude of ways. To compensate for any effect of caspase inhibition on early virus propagation, we began our treatment on day 2 pi and found no significant difference in viral titers on day 3 pi, the peak of viral replication in this model (36). Mice treated with the caspase-8 inhibitor showed reduced weight loss and improved TR-AM survival after IAV infection. On day 7 pi, caspase-8 inhibition mitigated without completely preventing TR-AM loss, potentially through the abundance of death-inducing signals on that time point and an escape towards the necroptotic pathway (57).

Through a stochastic interrogation of TNFSF member expression, we found a significant upregulation of the TNFSF14 ligand in the lungs of infected mice. High levels of soluble TNFSF14 were also detected in the BALF of patients with severe virus-induced ARDS. Previous studies on severe viral pneumonia and sepsis have demonstrated a positive correlation between BALF/serum TNFSF14 and disease severity (20, 21, 58, 59). TNFSF14 can exert a variety of roles on cell survival, profile (re)programming, establishment of immune response and infection memory, depending on cell type, pathogen interaction, and receptor availability (15, 22, 60-62). TNFSF14 has been previously described as a determinant of macrophage survival, phenotype, and antibacterial properties (63-65), however, extensive studies regarding any association with post-influenza TR-AM death are lacking. In our study, elimination of the ligand either through genetic deletion or through specific antibody blockade led to the preservation of the TR-AM pool, improved survival, and a lower lung bacterial burden after a secondary pneumococcal hit. We identified neutrophils as the main leukocyte source of TNFSF14, which is in accordance with previously published data (66, 67). TNFSF14 has been shown to play an instrumental role in NK and T cell activation and expansion (16, 23, 60, 68) and DC maturation (69) and may thus serve as an intermediate between the acute and adaptive immune response within the infection context. Aberrant release due to dysregulated neutrophil activation could offer an alternative explanation for the abundant TNFSF14 presence upon severe infection. This is in accordance with literature, as high circulating or organ-specific TNFSF14 levels have been positively correlated with highly inflammatory states, both pathogen-driven and autoimmune (18, 20, 70-73), and blocking of the ligand was shown to limit inflammation and attenuate organ injury (74).

TNFRSF14 and LTβR, the two competitor TNFSF14 receptors, followed distinct kinetics in terms of transcriptional regulation and protein expression on TR-AM during the infection course. Out of the two receptors, LTβR was shown to have a stronger impact on TR-AM death, as *ltβr^-/-^* mice showed an attenuated TR-AM loss after IAV infection, as opposed to *tnfrsf14^-/-^* mice, and intrapulmonary transfer of *ltβr^-/-^* TR-AM into wt mice significantly improved survival after co-infection, compared to transfer of *tnfrsf14^-/-^* or wt TR-AM. Given that more prominent TR-AM preservation was observed in *tnfsf14^-/-^* mice compared to *ltβr^-/-^* mice, we hypothesize that cell death could also be initiated through ligation of TNFSF14 to TNFRSF14, potentially through activation of a co-receptor, as TNFRSF14 lacks a pro-death domain, unlike other receptor members of the TNFSF, such as TNFRSF1 (15, 75). It should be noted, however, that the engagement of LTβR by TNFSF14 on monocytes is not merely confined to apoptosis induction. Transforming growth factor-beta (TGF-β) is secreted upon crosslinking, which has been shown to drive an immunoparalysis state in the aftermath of infection, further enhancing secondary infection susceptibility (76). LTβR-TNFSF14 interaction on the endothelium alters microvasculature structure, which can in turn favor the recruitment of immune cells (77). The intricate nature of TNFSF14-TNFRSF14/LTβR interactions thus points at a multitude of potential roles for the signaling axis in the context of influenza, besides TR-AM depletion. Nevertheless, with clinical trials in the context of virus-induced pneumonia and systemic inflammation already revealing beneficial safety profiles (59, 78, 79), therapeutic interventions disrupting TNFSF14-initiated intercellular pathways to preserve TR-AM function appear as promising novel approaches for improving host defense in the context of IAV pneumonia.

## Methods

### Sex as a biological variable

Sex was not considered as a biological variable for patient samples. Both male and female mice were used for all studies.

### Mice

Wt C57BL/6 mice were purchased from Charles River Laboratories. *Tnfsf14^-/-^* (80), *tnfrsf14^-/-^* (81), and *ltβr^-/-^* (82) mice were a gift from Prof. Klaus Pfeffer (Heinrich Heine University Düsseldorf, Düsseldorf, Germany). *Tnfsf10^-/-^* (83) mice were obtained from AMGen. All mice were bred under specific-pathogen-free conditions (SPF).

### In vivo infection

For *in vivo* IAV infection experiments, mice were o.t. inoculated with 250-1000ffu of A/Puerto Rico/8/1934 (PR8, H1N1) influenza virus. Control groups were inoculated with sterile PBS^-/-^. For co-infection experiments, mice were i.n. infected with 20cfu Spn [serotype 3, strain PN36 (NCTC 7978), provided by the group of M. Witzenrath, Department of Infectious Diseases and Pulmonary Medicine, Charité, University Medicine Berlin, Berlin, Germany] 7 days after IAV infection.

### In vivo treatment

For apoptosis inhibition, wt mice were infected with 500ffu or 1000ffu IAV and treated with s.c. injections of 10mg/kg of a specific caspase-8 inhibitor (Z-IETD-FMK, R&D Systems), or a DMSO control in a total volume of 60μl. For day 3 experiments, treatment involved a single injection on day 2 pi, whereas daily injections were applied from day 2-6 pi for analysis on day 7 pi. Neutrophil depletion was performed through the i.p. application of 200µg anti-1A8 antibody (*InVivo*Plus anti-mouse Ly6G, Cat BP0075-1, BioXCell) or an anti-2A3 isotype control (*InVivo*Plus^TM^ rat IgG2a isotype control, anti-trinitrophenol, Cat BP0089, BioXCell) diluted in 200µl sterile PBS in IAV-infected mice on days -1, 1, 3, and 5 pi. TNFSF14 neutralization was achieved with a mouse anti-mouse LIGHT blocking antibody (clone 3D11, isotype control mouse IgG2b, clone 27-35, BioLegend), kindly provided by Prof. José Ignacio Rodríguez Barbosa and Prof. Maria-Luisa del Rio (INBIOMIC, University of León, León, Spain). A single i.p. injection of 500µg of antibody or isotype control in 200µl PBS was performed two days after IAV infection. For *in vivo* treatment with rTNFSF14, 10µg of carrier-free mouse rTNFSF14 (Cat. 1794-LT, R&D Systems) diluted in 30µl PBS^-/-^ were o.t. administered to IAV-infected mice on days 1 and 2 pi for analysis on day 3 pi.

### Adoptive transfer of tissue-resident alveolar macrophages (TR-AM)

Murine TR-AM were obtained from the BALF of naïve wt, *tnfrsf14^-/-^,* and *ltβr^-/-^* mice, as previously described (34). Adoptive transfer was performed on day 3 after infection of wt mice with 250ffu IAV. Secondary pneumococcal infection was performed four days later.

### Clodronate depletion and co-infection

Two days prior to IAV infection, 70μl clodronate or control PBS liposomes (Liposoma, Cat. CP-005-005) were administered to wt and *tnfsf14^-/-^* mice. Pneumococcal infection was performed seven days after IAV infection. *Flow cytometry and FACS.* Multicolor flow cytometry analysis and FACS were performed on an LSR Fortessa and a BD FACSAria III cell sorter using the FlowJoTM v10.7 software (BD Biosciences), as previously described (34). Cells obtained from the BALF and homogenized lung tissue of naïve and infected animals were stained with 7-aminoactinomycin D (7-AAD, Cat. 420404, BioLegend) or Sytox^TM^ Blue Dead Cell Stain (Cat. S34857, Invitrogen) for dead cell exclusion. Total cell numbers were determined on a NucleoCounter® NC-3000TM (ChemoMetec). Primary antibodies included goat anti-mouse influenza A Virus (clone ab20841, abcam), rat anti-mouse CD45 APC/Cy7, PE/Cy7 (clone 30-F11, BioLegend), Ly-6G/Ly-6C (Gr-1) PE/Cy7 (clone RB6-8C5), SiglecF PE, BV421 (clone E50-2440, BD Biosciences), CD11c FITC, PE/Cy7 (clone N418, BioLegend), Ly-6G APC (clone 1A8, BioLegend), CD11b FITC, Pacific Blue (clone M1/70, BioLegend), CD3e FITC (clone 500A2, BioLegend), CD4 PE (clone GK1.5, BioLegend), CD8a PE/Cy7 (clone 53-6.7, BioLegend), CD19 PE (clone 6D5, BioLegend), CD206 APC (clone C068C2, BioLegend), CD24 PE/Cy7 (clone M1/69, BioLegend), CD64 (FcγRI) Pacific Blue (clone X54-5/7.1, BioLegend), CD326 (Ep-CAM) APC/Cy7 (clone G8.8, BioLegend), CD31 Alexa Fluor 488 (clone 390, BioLegend), and mouse anti-mouse NK1.1 APC (clone PK136, BD Biosciences). Secondary antibodies included rabbit anti-goat IgG (H+L) cross-adsorbed secondary antibody Alexa Fluor-647 (clone A-21446, Invitrogen) and goat anti-rabbit IgG (H+L) secondary antibody FITC (clone 31635, Invitrogen). Flow-sorted cells were resuspended in 350µl RLT^TM^ buffer (Qiagen) for further gene expression analyses.

### Immunohistochemistry

Lungs of IAV- and PBS-infected mice were fixed for 24h in 4% paraformaldehyde, embedded in paraffin, and sliced in 3-5 µm thick sections. Antigen retrieval was performed through protease addition (0,1g protease from Streptomyces griseus, Pan Reac AppliChem, ITW Reagents, activity: 10474,48 U/mg) in 100 ml PBS (10’, 37°C). Blocking serum [PBS, Roti-ImmunoBlock (Roth) and goat normal serum] was applied for 30min at RT and the primary antibody (TNFSF14 ab203578, goat-anti-rabbit, abcam) was applied overnight at 4°C. The next day, slides were incubated in the secondary antibody [goat-anti rabbit IgG (H+L), biotinylated, BA-1000, Vector Laboratories)]. Images were obtained at the Olympus BX 41 microscope using the cellSens imaging software (Olympus Corporation ver. 1.18) and were edited with Photoshop CS5 (64 bit). TNFSF14 expression was determined in a semi-quantitative manner, based on the frequency and intensity of positive signal in the lung interstitium, airspaces, subpleural regions, and bronchial epithelium. A quantification system was developed, ranging from 0 -no signal present-to 4, when TNFSF14 signal was vividly present in the examined tissue.

### TR-AM isolation and cell culture for ex vivo treatment

Following BALF extraction from naïve mice, cells were resuspended in full TR-AM medium (RPMI-1640/2% fetal bovine serum (FBS)/2.5% HEPES/1% L-glutamine/1% penicillin/streptomycin). TR-AM were seeded at a density of 10-50,000 cells/well on a 96-well plate.

### Colorimetric viability assay for ex vivo treated TR-AM

Primary TR-AM isolated from the BALF of naïve wt mice were treated with 0.1ml TR-AM medium containing 10% UV-inactivated, cell- and virus-free BALF. For caspase inhibition experiments, cells were pre-incubated in 50µM of a specific caspase-3 (Z-DEVD-FMK, R&D Systems) or caspase-8 inhibitor (Z-IETD-FMK, R&D Systems) for 3h prior to BALF treatment. Viability was assessed by a colorimetric assay (Cell Counting Kit-8, Sigma Aldrich), as per the manufactureŕs instructions. Absorbance was measured in an iMark microplate reader (Bio-Rad).

### Caspase-3/7 and caspase-8 activity

Wt, *tnfrsf14^-/-^,* and *ltβr^-/-^* BALF TR-AM were treated with 0.1ml TR-AM medium containing 10% BALF from wt or *tnfsf14^-/-^* mice for 24h. For ligand blocking experiments, BALF had been previously incubated with 1µg/ml of the mouse anti-mouse TNFSF14 antibody or an isotype control for 1h at 4°C prior to BALF treatment. Lyophilized Caspase-Glo^®^ 3/7 or Caspase-Glo^®^ 8 substrate (Promega) was resuspended in 10ml luciferase-containing Caspase-Glo^®^ buffer (Promega) and added á 0.1ml/well to the cells. After 1h incubation at RT, cells were transferred to a black 96-well plate for luminescence detection using a 520/25 filter in an FLx800 fluorescence reader (BioTek Instruments).

### Phagocytosis assay

Wt and *tnfsf14^-/-^* TR-AM and BMDM were compared regarding their phagocytic capacity on day 7 pi, using the pHrodo^TM^ Green *Escherichia coli* BioParticles^TM^ phagocytosis kit for flow cytometry (Invitrogen) as per the manufactureŕs instructions. Flow cytometry analysis was performed on an LSR Fortessa, as described above.

### Spn load after in vivo and ex vivo infection

Two days after IAV infection and 48h after Spn infection, BALF, lungs, and spleens from co-infected mice were harvested and homogenized. A series of inoculum dilutions in NaCl was prepared for each sample in 1:10 steps. For each dilution step, 4 x 0.01ml inoculum were pipetted on a blood agar plate and stored at 37°C overnight. Total bacterial load was calculated by counting the average number of separately grown colonies, multiplied by 10^number of dilution step*100^ (= number of colonies in 1ml). Wt and *ltβr^-/-^* TR-AM were *ex vivo* infected with Spn at an MOI 1000 for 10min, after which cells were vigorously washed 5 times in ice-cold PBS to remove any extracellular bacteria and lysed in water. An inoculum dilution series was pipetted on blood agar plates as described above. Total colony count on the following day depicted phagocytosed bacteria.

### ELISA

Soluble TNFSF14 was measured in murine and human BALF samples using commercially available ELISA kits (murine TNFSF14: R&D Systems, cat. DY1794-05, detection limit 250pg/ml, human TNFSF14: R&D Systems, cat. DLIT00, detection limit 16.5pg/ml), according to the manufacturer’s instructions.

### Quantitative PCR

RNA isolation was performed with the RNeasy kit (Qiagen) and cDNA was synthetized according to the manufactureŕs instructions. Quantitative PCR (qPCR) was then performed with SYBR green I (InvitrogenTM) in the StepOnePlusTM Real-Time PCR System and in the QuantStudio™ 3 Real-Time PCR System (Applied BiosystemsTM). *Gapdh* and *RPS18* expression served as normalization controls. Data are presented as ΔΔCt [(Ct reference– Ct gene of interest)_inf_ -(Ct reference– Ct gene of interest)_control_] or fold change (2ΔΔCt). The following primers were used:

a. murine: *Gapdh* (forward primer, 5’-AGGTCGGTGTGAACGGATTTG-3’; reverse primer, 5’-GGGGTCGTTGATGGCAACA-3’), *Tnfsf14* (forward primer, 5’-ACTGCATCAACGTCTTGGAGA-3’; reverse primer, 5’-TGGCTCCTGTAAGATGTGCTG-3’), *Tnfrsf14* (forward primer, 5’-CAGGCCCCTACAGACAACAC-3’; reverse primer, 5’-ACTCGTCTCCCACAAGGAACT-3’), *Ltβr* (forward primer, 5’-CCCCTTATCGCATAGAAAACCAG-3’; reverse primer, 5’-TGCATACCGCAAAGACAAACT-3’)
b. human: *TNFSF14* (forward primer, 5’-GGTCTCTTGCTGTTGCTGATGG-3’; reverse primer, 5’-TTGACCTCGTGAGACCTTCGCT-3’), *RPS18* (forward primer, 5’-GCAGAATCCACGCCAGTACAAG-3’; reverse primer 5’-GCTTGTTGTCCAGACCATTGGC-3’).

### RT^2^ Profiler PCR Arrays

RT^2^ profiler PCR arrays (Qiagen) were used for pathway or group gene expression analysis of flow-sorted TR-AM. RNA isolation, genomic DNA elimination, reverse transcription, cDNA synthesis, and qPCR were performed according to the manufactureŕs instructions. In samples with low RNA amount (<1µg), a pre-amplification of cDNA targets preceded qPCR by addition of a PCR mastermix (RT^2^ PreAMP PCR Mastermix) and a species- and pathway-specific primer mix (PBM-063Z-RT^2^ PreAMP cDNA Synthesis Primer Mix for Mouse TNF Ligands and Receptors, PBM-212Z - RT² PreAMP cDNA Synthesis Primer Mix for Mouse Cell Death PathwayFinder^TM^) to the cDNA samples. PCR components were added to the 96-well plate format provided by the company (PAMM-063ZC-24-RT² Profiler™ PCR Array Mouse TNF Signaling Pathway, PAMM-212ZC-12-RT² Profiler™ PCR Array Mouse Cell Death PathwayFinder). Real-time PCR was performed in the StepOnePlusTM Real-Time PCR System and in the QuantStudio™ 3 Real-Time PCR System (Applied Biosystems^TM^). Data analysis was performed with the company’s web-based data analysis software (GeneGlobe Data Analysis Center, Qiagen).

### Single-cell RNA sequencing

To identify the main TNFSF14 source upon IAV infection, 500,000 live leukocytes (gated as Sytox-CD45+) from the homogenized single-cell lung suspensions of wt IAV-infected mice on day 3 and 7pi were sorted into 0.35μL 0.04% BSA/PBS^-/-^. Following viability control, 10,000 cells were loaded onto Chromium Chip B (10X Genomics). Gel beads in emulsion (GEM) generation, cDNA synthesis and amplification, and library preparation were performed with the Chromium Single Cell 3’ Reagent Kit v3.1 (10X Genomics), as per the manufacturer’s protocol. Indexed libraries were sequenced on an Illumina NextSeq2000. Prior to analysis, reads were aligned against the mouse genome (GRCm38.p6) and quantified using StarSolo (https://github.com/alexdobin/STAR). Analysis was conducted with the Scanpy software (https://github.com/theislab/scanpy). After quality filtering, raw cell counts were normalized to the median count over all cells and transformed into log space for variance stabilization. Principal component analysis (PCA) identified 14 and 11 components on days 3 and 7 pi, respectively. Uniform Manifold Approximation and Projection (UMAP) embedding was created to identify cell type clusters through Leiden clustering. Doublet analysis was conducted on day 7 pi using Scrublet (https://github.com/swolock/scrublet), leading to the removal of a doublet cluster (118 cells).

### Neutrophil isolation from human BALF samples

Neutrophil isolation from the BALF of patients with IAV ARDS and patients who underwent bronchoscopy for diagnostic purposes was performed with the MACSxpress® Whole Blood Neutrophil Isolation Kit (Cat. 130-104-434, Miltenyi), according to the manufactureŕs instructions.

### Statistics

Data are shown in scatterplots as single data points. Means ± SEM per group are indicated by bars and error bars. Statistical significance between two groups was calculated using student’s t-test. For comparison between more than two groups, significance was determined by 1-way ANOVA, 2-way ANOVA with Tukey’s posthoc test, or by Kruskal-Wallis test followed by Dunn’s post hoc comparison test. Survival curves were compared by log-rank (Mantel-Cox) test. p<0.05 was considered as significant. \**P* < 0.05; \*\**P* < 0.01; \*\*\**P* < 0.005. Graphs were prepared using GraphPad Prism (GraphPad Software version 10.2.3).

### Study approval

Animal experiments were approved by the regional authorities of the State of Hesse (Regierungsprasidium Giessen) and by the Institutional Ethics Committee at the IBioBA Institute. Use of human BALF samples was approved by the University of Giessen Ethics Committee, samples were provided by the biobank of the German Center for Lung Research (DZL). Written informed consent was received prior to sample use.

### Data availability

Detailed experimental methods, materials, and data supporting the findings of this study are available within the article and its supplemental material. Values for all data points in graphs are reported in the Supporting Data Values file.

## Author contributions

CM, CP, MRF, JB, ME designed and performed experiments, evaluated, and interpreted data. CM wrote the manuscript. AIVA and SH designed, performed, and interpreted scRNA-Seq experiments. IA performed imaging analysis. HS, SG, ML performed sequencing and bioinformatic analyses. JH and ADG performed immunohistochemistry analysis. KF provided genetically modified mice. MLDR and JIRB provided the neutralizing anti-TNFSF14 antibody (clone 3D11). IV performed bronchoscopy for the acquisition of human BALF samples. CP, SG, ML, ADG, IV, UM, and SH revised the manuscript. SH conceived the scientific question, designed experiments, interpreted data, and financed the study.

## Supporting information

Figure values with mean and SD

## Acknowledgements

This work was supported by the German Research Foundation (Deutsche Forschungsgemeinschaft, DFG) under following projects: KFO309 project P2/P9, project number 284237345; SFB-TR84 project number 114933180; SFB1021 project number 197785619; the German Center for Lung Research (Deutsches Zentrum für Lungenforschung, DZL), and the Cardiopulmonary Institute (CPI; DFG EXC 2026 project number 390649896). We would like to thank Martin Witzenrath and Birgit Gutbier for providing us with the S bacterial strain. We would like to thank Maria Groß, Stefanie Jarmer, Larissa Hamann, Florian Lück, Julia Stark, and Josephine Guth, for excellent technical support.

---

**Supplemental Figure 1.**
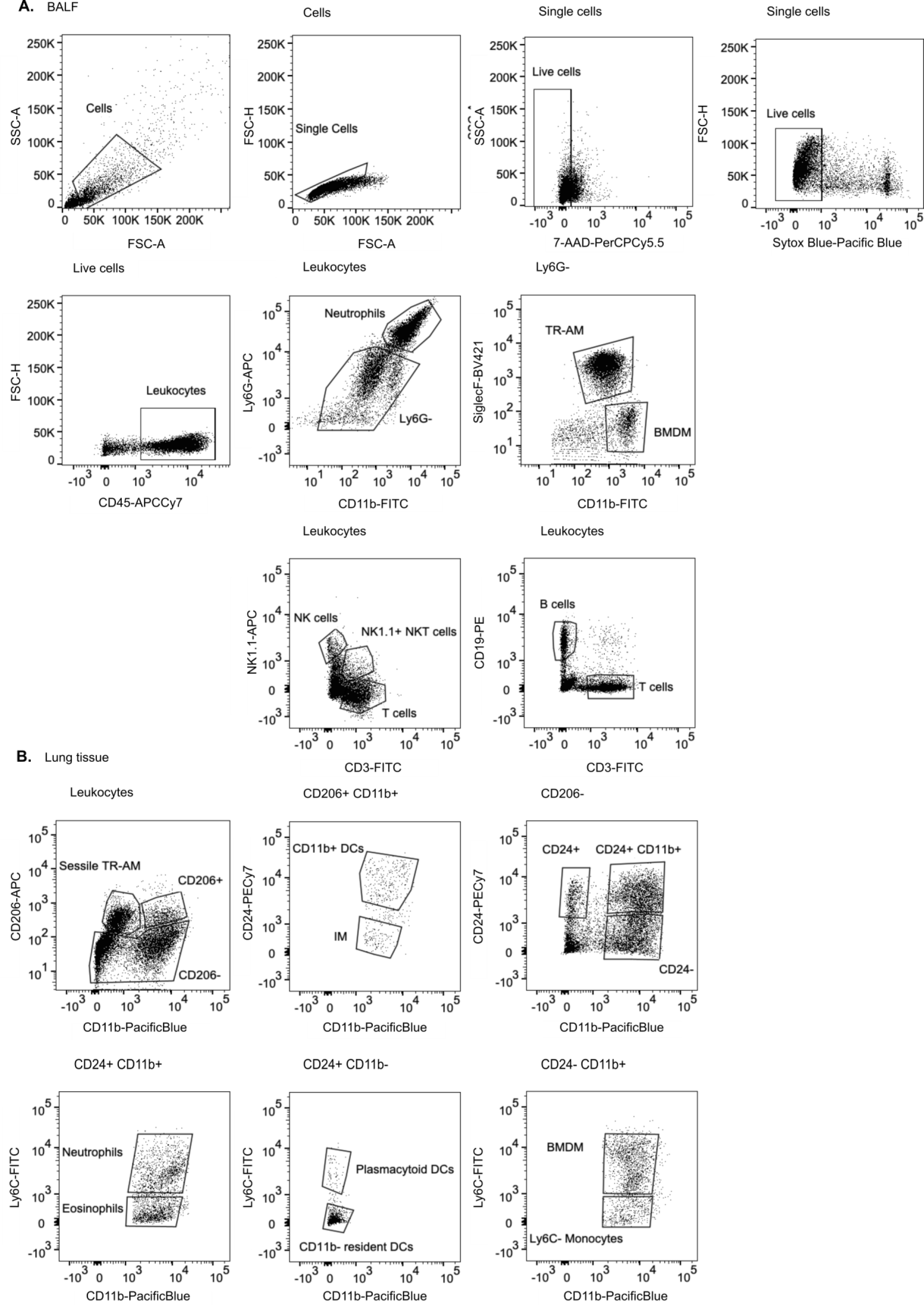
Gating strategy for the identification of different immune cell populations in murine BALF (A) and in the lung tissue (B) per flow cytometry analysis. Representative plots following multicolor staining of BALF and lung-tissue immune cells, as described in the ‘Methodś section. Gating strategies were set according to the appropriate isotype and fluorescence minus one (FMO) controls.

**Supplemental Figure 2.**
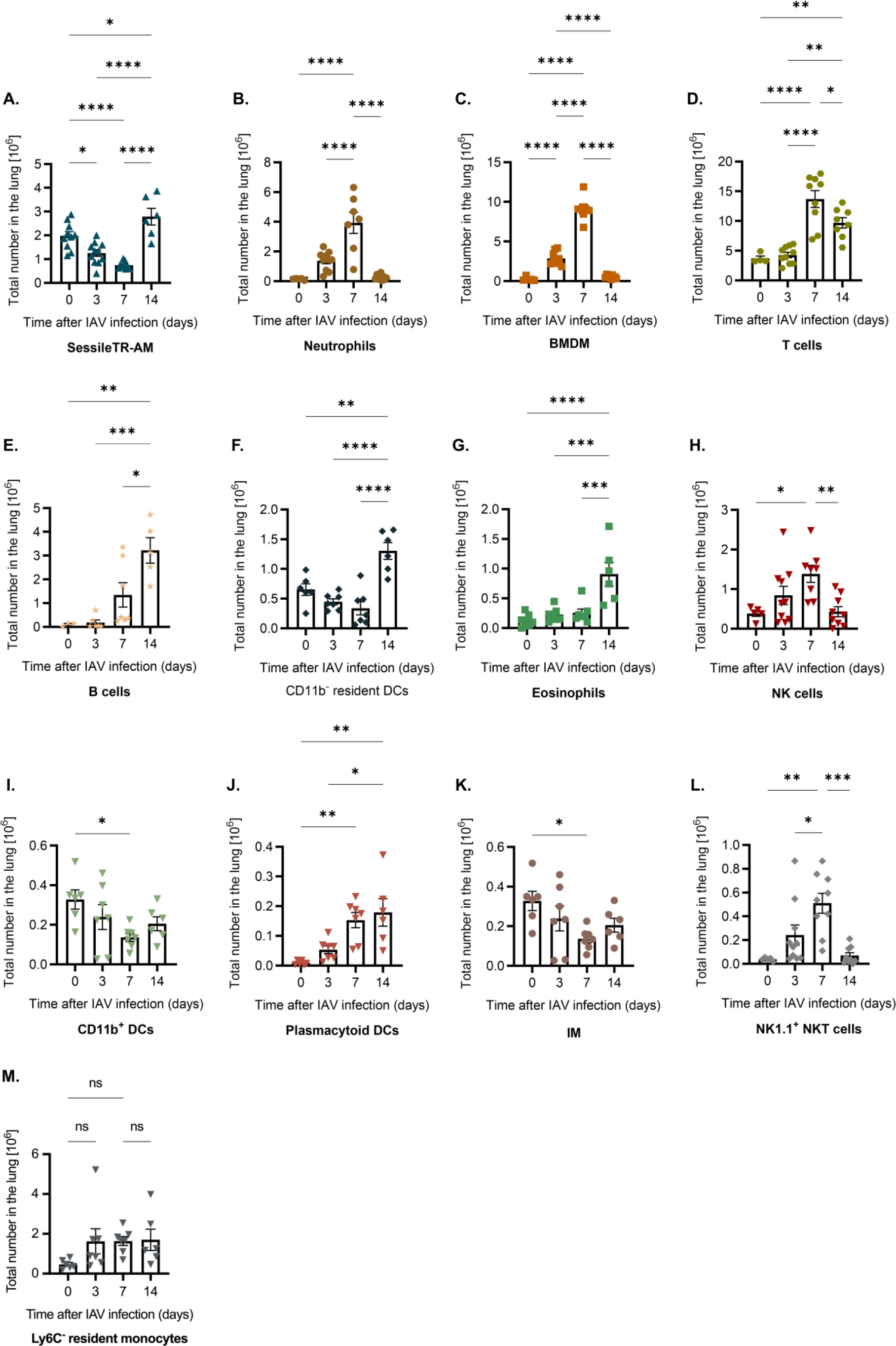
Immune cell kinetics in the lung tissue of IAV-infected mice over the infection course. Following BALF extraction, lungs of wild-type mice infected with 500ffu IAV were harvested at different time points. Immune cell populations were identified per flow cytometry analysis, including sessile TR-AM (n=6-11) **(A)**, neutrophils (n=5-10) **(B)**, BMDM (n=5-10) **(C)**, T cells (n=4-10) **(D)**, B cells (n=3-7) **(E)**, CD11b^-^ resident DCs, (n=6-7) **(F)**, eosinophils (n=6-7) **(G)**, NK cells (n=5-10) **(H)**, CD11b^+^ DCs (n=6-7) **(I)**, pDCs (n=6-7) **(J)**, IM (n=6-7) **(K)**, NK1.1^+^ NKT cells (n=5-10) **(L)**, and Ly6C^-^ resident monocytes (n=6-7) **(M)**. Data shown are pooled from five different experiments, mean ± SEM is depicted. Significance was determined by 1-way ANOVA with Tukey’s posthoc test; *p<0.05, **p<0.01, ***p <0.001, ****p <0.0001.

**Supplemental Figure 3.**
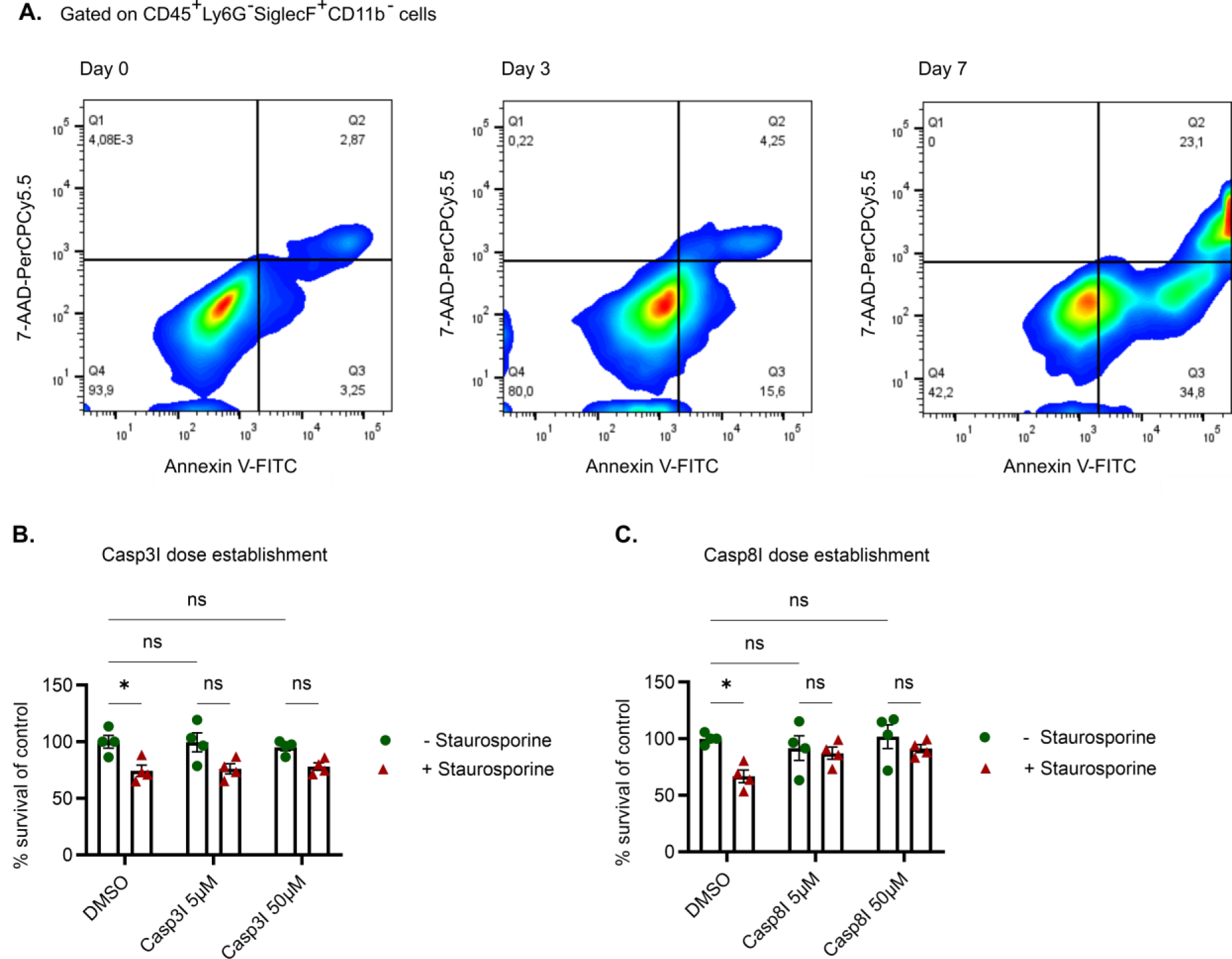
Gating strategy for the quantification of apoptotic TR-AM after IAV infection and establishment of *ex vivo* caspase inhibitor treatment. **(A)**Gating strategy for the quantification of apoptotic TR-AM in the BALF of IAV-infected mice on days 0, 3, and 7 pi. **(B-C)** TR-AM viability assessment following treatment with 0.5μM staurosporine and different doses of a caspase-3 **(B)** or a caspase-8 **(C)** inhibitor, values depicted as % survival of control. Non-treated cells set at 100%. Graphs represent means ± SEM, n=4 per condition. Significance was determined by 2-way ANOVA with Tukey’s posthoc test; *p<0.05.

**Supplemental Figure 4.**
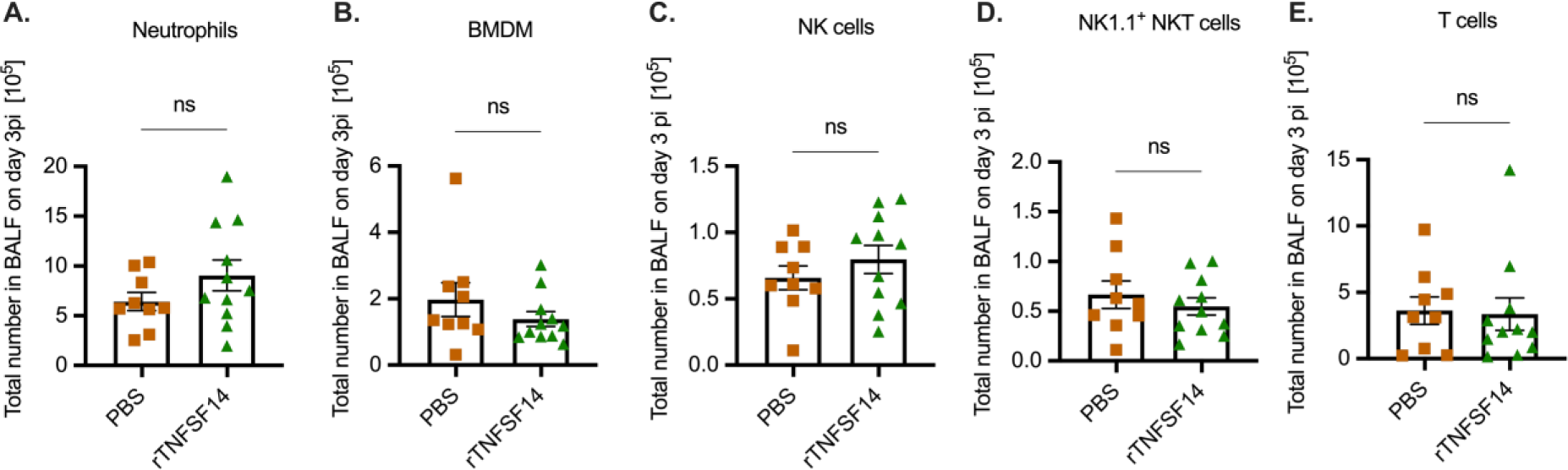
T**N**FSF14 **orchestrates post-influenza TR-AM death. (A-E)** BALF numbers of BMDM **(A)**, neutrophils **(B)**, NK cells **(C)**, NK1.1^+^ NKT cells **(D)**, and T cells **(E)** on day 3 pi after PBS or rTNFSF14 treatment of wt mice on days 1 and 2 pi. Graphs represent means ± SEM, n=6-8. Significance was determined by unpaired t-test; *p<0.05.

**Supplemental Figure 5.**
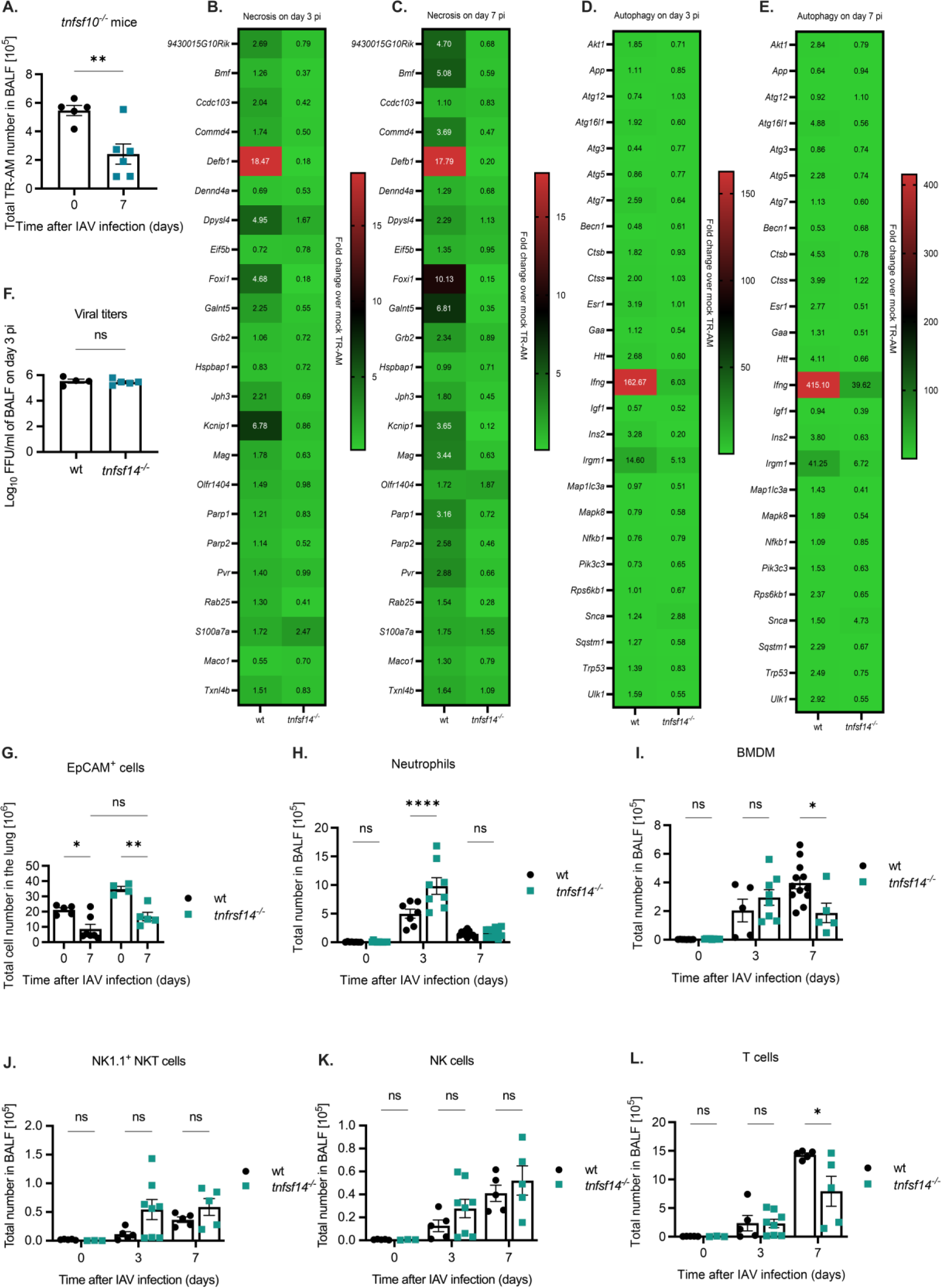
Post-influenza TR-AM death is attenuated in the absence of the TNFSF14 ligand. **(A)**Quantification of TR-AM numbers in the BALF of uninfected and IAV-infected *tnfsf10^-/-^* mice on day 7 pi (means ± SEM, n=5-6). **(B-E)** Heat maps depicting fold change of necrosis-**(B-C)** and autophagy-related **(D-E)** genes on day 3 pi **(B, D)** and day 7 pi **(C, E)** over mock-infected wt and *tnfsf14^-/-^* TR-AM, n=3-5, as determined per gene array analysis. **(F)** Viral titers in the BALF of IAV-infected wt and *tnfsf14^-/-^* mice on day 3 pi (means ± SEM, n=4 animals per group). **(G)** Epithelial (EpCAM^+^) cells in the lungs of wt and *tnfsf14^-/-^* mice during IAV infection course (means ± SEM, n=4-7). **(H-L)** Immune cell populations in the BALF of wt and *tnfsf14^-/-^* mice over the early course of IAV infection, including neutrophils **(H)**, BMDM **(I)**, NK1.1^+^ NKT cells **(J)**, NK **(K)**, and T cells **(L)**. Graphs represent means ± SEM, n=5-15. Significance was determined by unpaired t-test and 1-way ANOVA with Tukey’s posthoc test; *p<0.05, **p<0.01, ***p <0.001, ****p <0.0001.

**Supplemental Figure 6.**
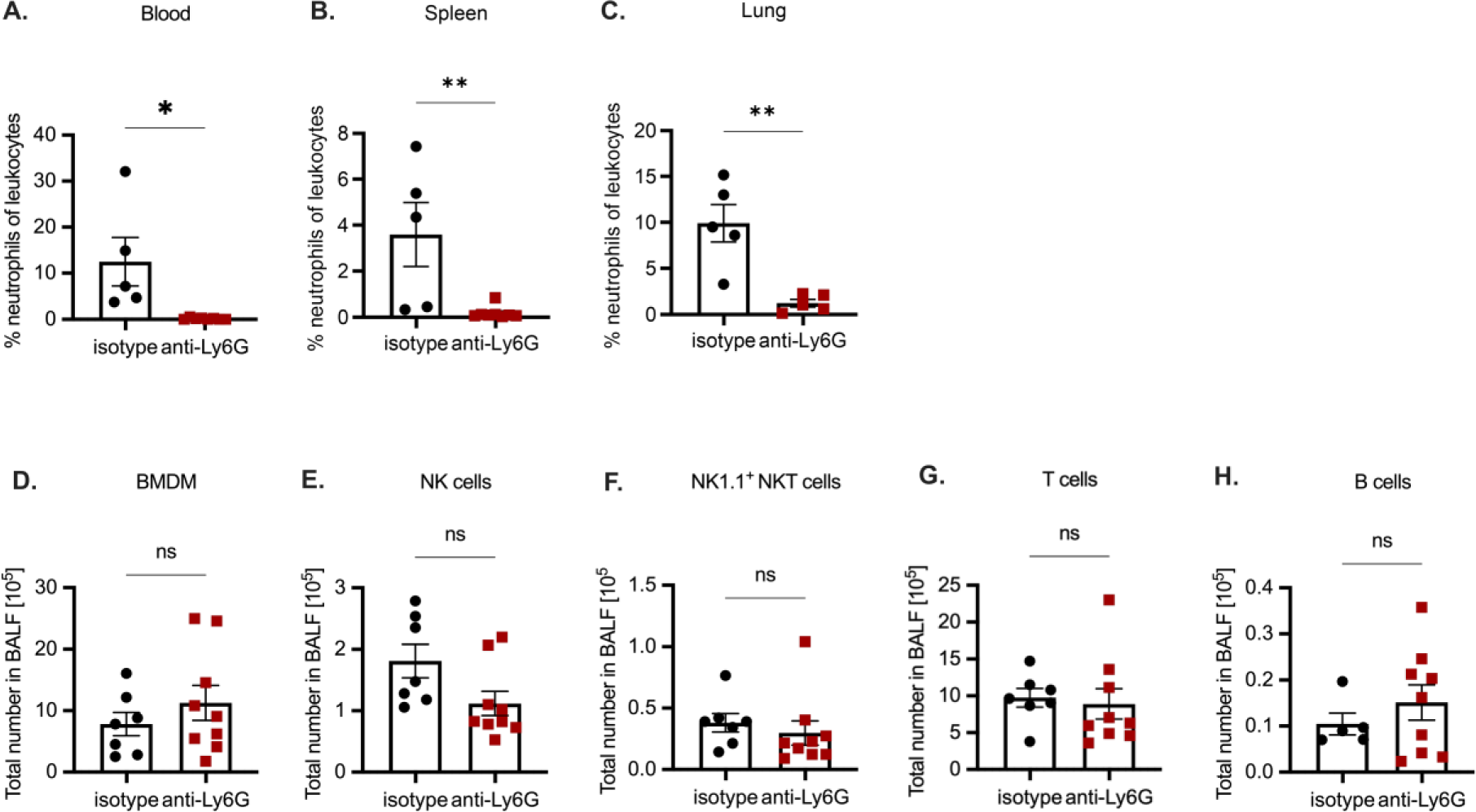
Neutrophil depletion attenuates post-influenza TR-AM loss. (A-C) Neutrophils depicted as percentage of total leukocyte numbers in the blood **(A)**, spleen **(B)**, and lungs **(C)** of IAV-infected animals after neutrophil depletion(means ± SEM, n=5-8). **(D-H)** Quantification of immune populations in the BALF of IAV-infected after anti-Ly6G or isotype control treatment, including BMDM, **(D)**, NK cells **(E)**, NK1.1^+^ NKT cells **(F)**, T cells **(G)**, and B cells **(H)**. Graphs represent means ± SEM, n=7-9. Significance was determined by unpaired t-test for two groups; *p<0.05, **p<0.01.

**Supplemental Figure 7.**
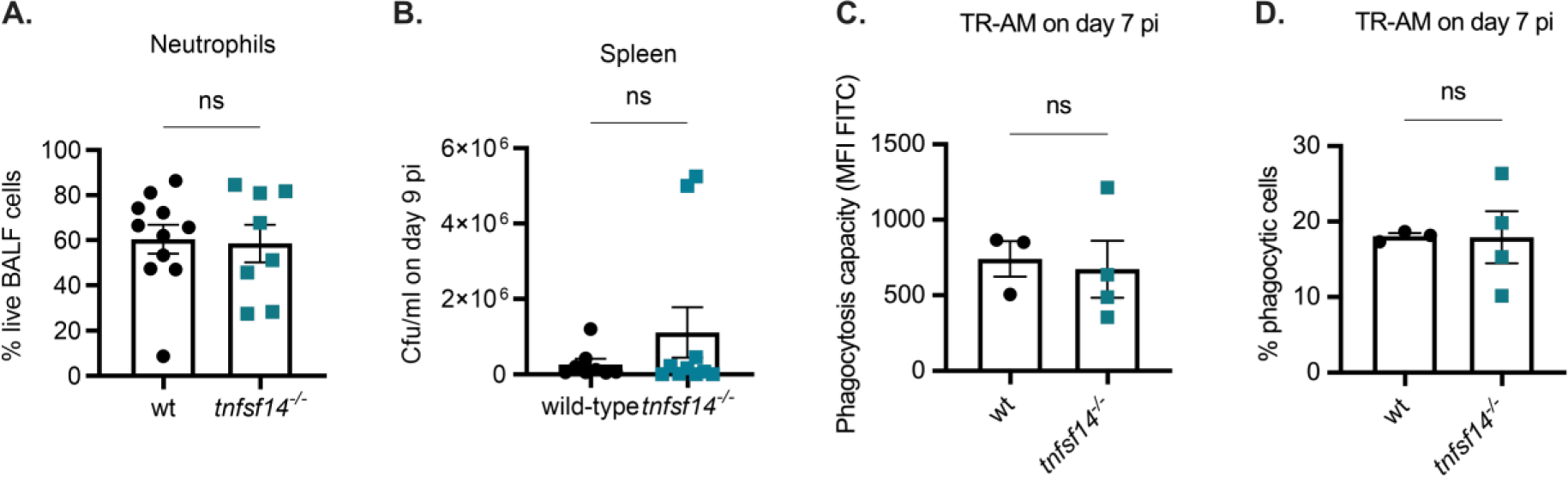
Loss of TNFSF14 improves outcome after IAV/Spn co-infection. **(A)**Neutrophils depicted as percentage of total live cells in the BALF of wt and *tnfsf14^-/-^* mice on day 9 pi after IAV infection and 48h after Spn infection (means ± SEM, n=8-10). **(B)** Spn burden in the spleens of wt and *tnfsf14^-/-^* mice 9 days after IAV infection and 48h after Spn infection (means ± SEM, n=7-9). **(C-D)** Phagocytosis capacity of TR-AM depicted as MFI FITC **(C)** and percentage of phagocytic TR-AM (% FITC+) out of total BALF TR-AM **(D)** on day 9 pi (means ± SEM, n=3-4). Significance was determined by unpaired t-test.

